# Extending the osmophobic effect to protein side chains with a physical transfer model across osmolyte classes

**DOI:** 10.64898/2026.06.17.732849

**Authors:** Ander F. Pereira, Jéssica O. Araújo, Wilson A. Tárraga, Leandro Martínez

## Abstract

Understanding the role of the protein backbone and side chains on cosolvent-induced stabilization is essential for a molecular picture of osmolyte action. The prevailing view holds that protecting osmolytes stabilize proteins mainly through unfavorable backbone interactions - the osmophobic effect - with side chains playing a minor or opposing role. By revisiting the decomposition of amino acid transfer free energies to properly account for mutual shielding between backbone and side chains, we derive a transfer model consistent with experimental denaturation *m*-values for both urea and protecting osmolytes - a feat neither existing model achieves alone. Backbone accessibility is mechanism-dependent: geometric for excluded cosolvents, complete for binders with specific cosolvent-backbone interactions. For strong protecting osmolytes - TMAO, sarcosine, sucrose, trehalose, and sorbitol - both backbone and side chains contribute favorably to stabilization, with side-chain contributions comparable or even greater than the backbone’s. For urea, accounting for the preferential binding and directional nature of urea-backbone hydrogen bonding - which makes the backbone accessible regardless of side-chain shielding - recovers the known balanced backbone and side-chain contributions to denaturation. Weaker protectants such as proline, betaine, and glycerol show competing backbone and side-chain effects that partially cancel. These results extend the osmophobic effect to side chains and yield a three-tier classification of osmolyte action: cooperative stabilization, cooperative destabilization, and competing backbone/side-chain contributions. The model’s greater sensitivity to side-chain composition offers avenues for experimental validation and for protein engineering.

## 1 Introduction

Naturally occurring osmolytes are small, soluble organic molecules that cells accumulate in response to osmotic, thermal, and desiccation stress [1, 2, 3, 4]. They are found across the tree of life — from bacteria in high-salinity environments to deep-sea cartilaginous fish, which counteract the denaturing effects of urea with high intracellular concentrations of trimethylamine *N* -oxide (TMAO) [5, 6], and to tardigrades and resurrection plants, which accumulate trehalose during desiccation [1]. Protecting osmolytes such as TMAO, sucrose, trehalose, sorbitol, sarcosine, and glycine betaine stabilize proteins at high concentrations without perturbing their function, making them indispensable for proteostasis [7, 8] and widely used as excipients against degradation of protein biotherapeutics [9]. Chemical denaturants such as urea and guanidinium chloride act in the opposite direction, unfolding proteins at moderate concentrations, and are routine tools to study folding equilibria.

The molecular basis of osmolyte action is rooted in their preferential interaction with — or exclusion from — the protein surface [10, 11]. Protecting osmolytes are excluded from the protein hydration shell, making the transfer of protein surface area from water to cosolvent unfavorable; because denaturation exposes more surface, the native state is selectively stabilized [7, 8]. Denaturants instead accumulate at the protein surface, favoring the exposure of buried groups and thus the unfolded state [10]. Tanford formalized this through the transfer free energy (TFE) — the free energy cost of moving a protein from water to a cosolvent solution — and showed that it decomposes into additive contributions from side chains and backbone units [12, 11]. The experimental denaturation *m*-value, which measures how strongly a cosolvent shifts the folding equilibrium, is then proportional to the TFE difference between native and denatured states, and can be predicted from the amino acid composition and accessible surface areas [12]. This group-additive picture has been extensively validated for many proteins and cosolvents, and recent tools make these calculations accessible at proteome scale [9, 13, 41]. A central question, however, is how the stabilizing (or destabilizing) effect is partitioned between the peptide backbone and the side chains — and whether any universal account of this partition is consistent with experimental data across all osmolyte classes. This question motivates the present work.

Here we first show that the two models currently in use display concurrent limitations that prevent their applicability to all classes of cosolvents, protectants and denaturants. We then propose a new model construction based on first principles, and demonstrate that accounting for the specificity of osmolyte-backbone interactions reconciles the predictions for denaturants and protectants. The new model redistributes backbone and side-chain contributions for protecting osmolytes, implying a fundamentally different biophysical interpretation of the osmophobic effect. Predicting a greater sensitivity of transfer free energies to amino acid composition, it can be experimentally tested and has implications for protein design.

### 1.1 The Established model

The *Established* additive model for estimating protein denaturation *m*-values was proposed by Tanford and developed for practical utility in particular by Bolen and co-workers [12]. It is underpinned by two kinds of experimental data: (1) the transfer free energies of glycine polypeptides of increasing length, or of cyclic diglycine, giving the transfer free energy of a backbone unit, TFE^*bb*^; and (2) the transfer free energies of the complete amino acids, TFE_*aa*_.

Using TFE_*Gly*_ as a model for a backbone with terminal groups, the transfer free energy of the side chain, *sc*, of another amino acid *aa* is obtained as

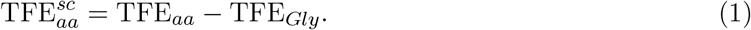

The TFE_*aa*_, including that of glycine, are obtained from the solubilities of the amino acids in water, 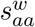, and in the target cosolvent solution, 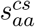, by

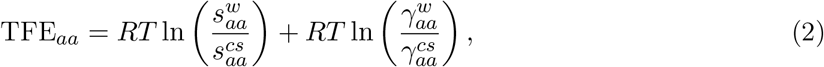

where 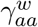 and 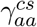 are the activity coefficients of the amino acids in each solution and 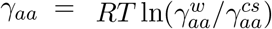. Solubilities measured for most natural amino acids in water and in various osmolyte solutions yield the *apparent* transfer free energies, 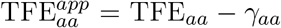. The side-chain contribution is then

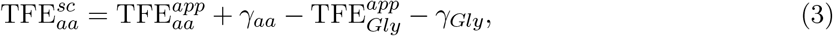

where *γ*_*aa*_ is found to be small, except for glycine in urea. By defining 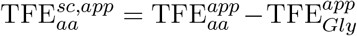 and using *γ*_*aa*_ = 0, the transfer free energy of a protein from water to a cosolvent can be obtained assuming group additivity:

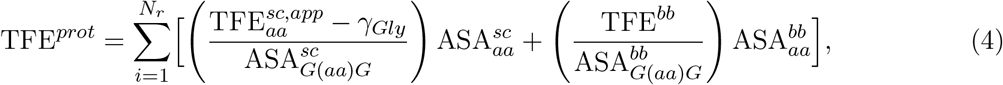

where *N*_*r*_ is the number of residues and the terms in parentheses are the transfer free energy per unit of surface area for the side chain or backbone of each amino acid type. The denominators were crucially computed from the solvent accessibility of the groups in *G*(*aa*)*G* tripeptides, representing fully solvent-exposed denatured states [12, 14, 11]. We revisit these accessible surface areas below. Denaturation *m*-values follow from the difference in transfer free energies between the native state and the denatured model of Creamer et al. [14]

The *Established* model, as described so far, successfully predicted transfer free energies and denaturation *m*-values across a variety of protecting osmolytes, roughly within experimental uncertainties [12, 15]. An inconsistency remained for urea, the most representative chemical denaturant, for which predictions were more negative than experimental values (Figure 1A). Bolen and co-workers [17] soon recognized that, specifically for urea, the activity correction for glycine could not be neglected in the estimation of the side-chain apparent transfer free energies (eq 4). Accounting for the non-ideality of glycine gave good agreement with experiment (Figure 1B), where TFE^*∗*^ denotes the glycine-activity-corrected transfer free energies [15]. The essential constituents of additive transfer models appeared to be settled.

**Figure 1:**
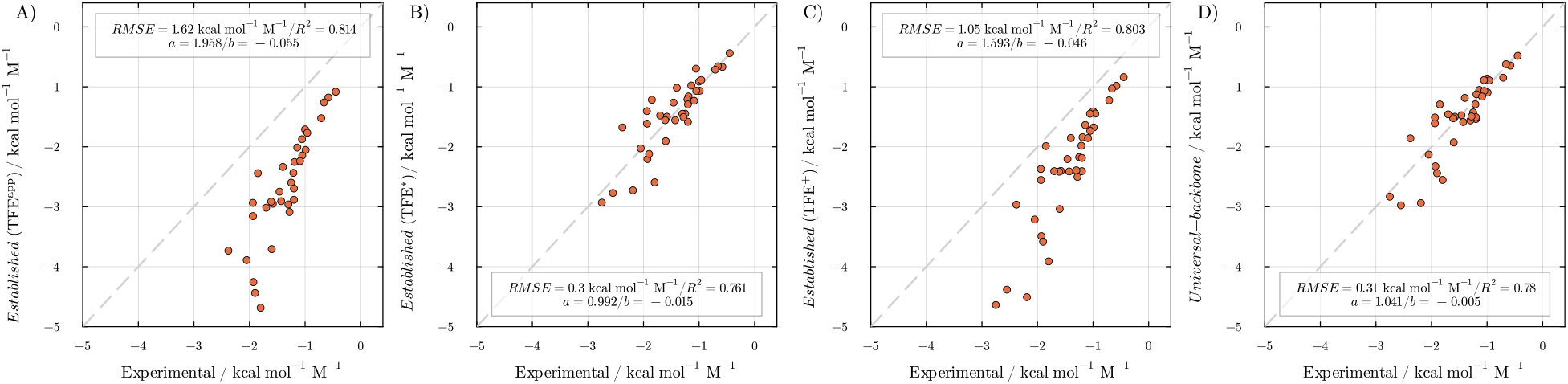
Experimental and predicted denaturation *m*-values in urea. A) *Established* model with apparent transfer free energies (TFE^*app*^). B) *Established* model with the incorrect glycine-activity parameter (TFE^*∗*^) [15]. C) *Established* model with the correct glycine-activity parameter (TFE^+^) [16]. D) *Universal-backbone* model with the correct glycine-activity parameter [16].

### 1.2 The Universal-backbone model

In 2013, this apparent consensus was broken. Moeser and Horinek found a concentration-unit conversion error in the non-ideality correction of the *Established* model, showing that a fortunate (or unfortunate) error cancellation had led to the good predictions for urea. When the proper glycine-activity correction is applied, the model again predicts a stronger denaturing role for urea than observed (Figure 1C), where TFE^+^ denotes the corrected 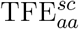 [16].

They noted that the inconsistency could be resolved by a subtle but important modification. In eq 4, the 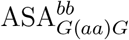 parameter is the solvent accessibility of the backbone of amino acid *aa* in a highly exposed tripeptide; it therefore carries the protection that the side chain of *aa* confers on the backbone and differs for each amino acid type, whereas TFE^*bb*^ is fixed for each cosolvent. They argued that the ASA of an exposed backbone does not depend on the amino acid type, so the denominator should be 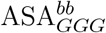 — the accessible surface area of the glycine backbone — which is chemically consistent, since all amino acids share the same backbone unit. Using amino-acid-specific ASAs overestimated the backbone contributions (negative for urea) because the transfer free energy per unit area was larger for all residues except glycine, so that with the proper glycine-activity correction the predicted *m*-values were again too negative (Figure 1C).

The model in which 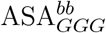 replaced 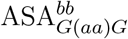 in eq 4 was named the “universal backbone” model. By reducing the (negative) backbone contributions with the greater ASA denominator, its *m*-value estimates for urea became again consistent with experiment (Figure 1D). The *Universal-backbone* model was important for a fundamental reinterpretation of backbone vs. side-chain contributions: while the *Established* model predicted a dominant role for the backbone, the *Universal-backbone* model predicted balanced contributions of side chains and backbones (Supplementary Figure S1), supported by simulations of Moeser and Horinek [16] and others [18, 19]. Once again, the fundamental structure of additive transfer models appeared to find a solid ground.

Balanced contributions are also supported by a series of additive transfer models proposed by Record and co-workers, parameterized from experimental data on the interaction of multiple small compounds with the target osmolyte [21, 22]. By fitting predictions grounded on atom-type decompositions (the Solute Partitioning Model) to the observed TFE, these models are more general and can be extrapolated to systems of greater structural complexity [23], such as protein structures, oligonucleotides, and macromolecular oligomers. We implemented the Record model with the available parameterizations for urea and other osmolytes (in PDBTools.jl [41] version 3.39.0), and computed the previously unreported implications of these models for the backbone vs. side-chain balance. For urea, the predictions agree qualitatively with the *Universal-backbone* model (Supplementary Information Section S3), and the key role of side-chain elements is recognized for other cosolvents [21, 22]. However, experimental data for training and validation are insufficient for a definitive analysis of backbone and side-chain roles for protecting osmolytes (Supplementary Information Sections S4 to S6).

## 2. Model Conflicts and Resolution

### 2.1 The tension between the models

The *Established* model remained the most widely used framework for estimating transfer free energies [9], despite the inconsistency found by Moeser and Horinek. Several reasons support its continued use: the total *m*-values predicted for urea with TFE^*∗*^ are as accurate as those of the *Universal-backbone* model; it predicts with comparable accuracy the total *m*-values for eight protecting osmolytes (TMAO, betaine, sarcosine, glycerol, sorbitol, proline, sucrose, and trehalose), even though the experimental data for each are sparser than for urea; and a practical web server has long been available for all these cosolvents [12]. Therefore, as long as the backbone-vs-side-chain partition is not the focus, its use appears justified.

The absence of flexible, general implementations of these models, including the alternative Record models, had nevertheless concealed a fundamental inconsistency. Because the *Universal-backbone* model reduces the backbone contributions, it must also reduce the total predicted *m*-values for any cosolvent with positive TFE^*bb*^. For urea this reduction is compensated by the glycine-activity correction, but for protecting osmolytes no analogous correction is available — glycine-activity corrections are expected to be small for these compounds — and the reduced backbone contributions cannot be offset. For sorbitol and glycerol, for example, Gekko et al. [20] estimate *γ*_*Gly*_ to be *∼* 0–2 and *∼ −*4 cal mol^*−*1^ M^*−*1^, much smaller than for urea. These small corrections are supported by the *Established* model predictions working in their absence, although independent experimental validation would be a valuable test.

Thus, the *Universal-backbone* model provides a physically sound decomposition of backbone and side-chain contributions to urea denaturation, consistent with simulations and with a chemically justified per-ASA backbone transfer free energy, yet it systematically underestimates the protective role of all protecting osmolytes (Figure 2). The *Established* model, conversely, correctly captures the total *m*-values for protecting osmolytes despite resting on a physically questionable partition, and gives effectively wrong predictions for urea when the correct glycine-activity correction is applied (Figure 1C). These contradictions are irreconcilable within the two frameworks: both rest on partially fallible physical assumptions concerning the detailed balance of group contributions.

**Figure 2:**
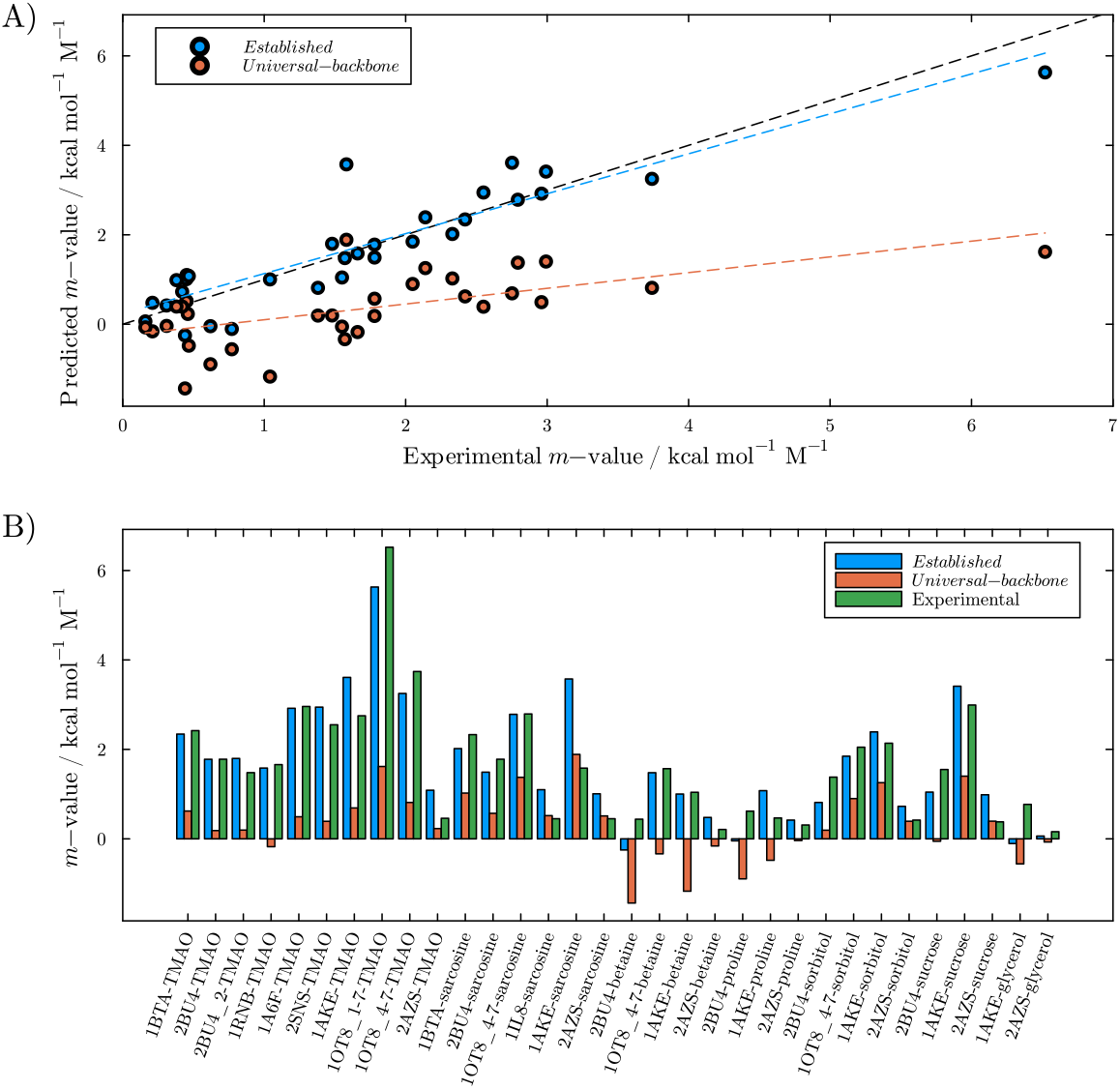
*Established* and *Universal-backbone* model predictions for protecting osmolytes. The *Established* predictions align with experimental data, but the *Universal-backbone* model underestimates the protective role systematically.

### 2.2 The Accessibility model

In view of the contradictory results each model provides for different osmolyte classes, we return to first principles. The new model shares features with both the *Established* and *Universal-backbone* models, but only as natural consequences of its physically motivated construction. It allows us to understand precisely where and why each previous model fails, and provides a unified interpretation of cosolvent effects across all osmolyte classes.

We start by revisiting the composition of the transfer free energy of an amino acid, which according to eq 1 should be TFE_*aa*_ = TFE^*sc*^ + TFE_*Gly*_. We instead decompose TFE_*aa*_ into side-chain, backbone, and terminal groups, as

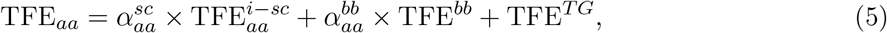

where TFE^*T G*^ is the transfer free energy of the terminal (capping) groups. Crucially, TFE^*i*-*sc*^ and TFE^*bb*^ are the transfer free energies of the *isolated* side chain and backbone: non-physical constructs in which the side chain is detached from the backbone, and the backbone is taken in the absence of any side chain. This adheres to strict group additivity and introduces the accessibility factors 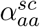 and 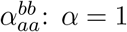 denotes a group fully accessible to the cosolvent and *α* = 0 a group fully shielded.

The TFE^*T G*^ term, which has no experimental counterpart, can be eliminated by noting that TFE_*Gly*_ = TFE^*bb*^ + TFE^*TG*^, since glycine consists of a backbone and terminal fragments only. Substituting into eq 5 and rearranging to obtain TFE^*i*-*sc*^ gives

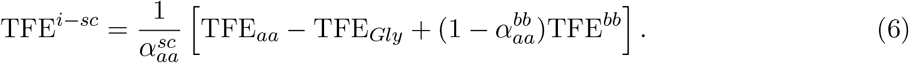

With 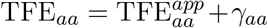 for any amino acid type (including TFE_*Gly*_) and 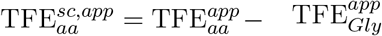, we obtain

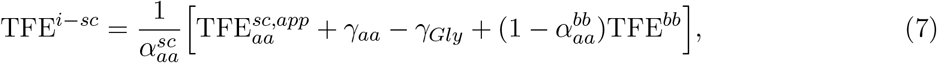

which is the transfer free energy of the isolated side chain of amino acid *aa*. The *γ*_*aa*_ term is assumed to be small (specific values for alanine in urea are available in the literature), and the thermodynamic data needed to evaluate TFE^*i*-*sc*^ are available from Auton and Bolen [12, 15]. It remains to specify the accessibility parameters.

#### 2.2.1 ASA based solvent accessibility

The natural way to express the accessibilities in eq 7 is through solvent-accessible surface area calculations. The ASAs computed by Creamer et al. [14] that parameterized the *Established* and *Universal-backbone* models cannot be used here, because Creamer defined the ASA of an exposed side chain as that in *G*(*aa*)*G* tripeptides, to model maximum solvent exposure in the denatured state. We need instead the non-physical ASA of the side chain isolated (detached) from the backbone, ASA^*i*-*sc*^, and the surface area of the side chain in the presence of the backbone, ASA^*sc*^, such that

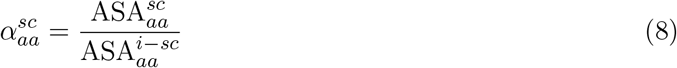

is the fraction of the side chain exposed to solvent in the presence of the backbone for a specific amino acid type. Similarly, the fraction of the backbone accessible to solvent when the side chain is present is

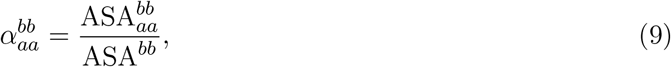

where 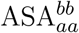is the accessible surface area of the backbone in the presence of the side chain, and ASA^*bb*^ is the area of the isolated backbone unit, independent of amino acid type and equal to 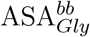, that of glycine. We computed these areas throughout the CATH S20 non-redundant protein database (see Methods); average values are given in Table 1.

**Table 1.** Average accessible surface areas (in Å^2^) and exposed fractions for isolated amino acids computed over the CATH S20 non-redundant protein database.

| AA | $ASA_{aa}^{sc}$ | $ASA_{aa}^{i-sc}$ | $ASA_{aa}^{bb}$ | $ASA^{bb}$ | $\alpha_{aa}^{sc}$ | $\alpha_{aa}^{bb}$ |
| --- | --- | --- | --- | --- | --- | --- |
| ALA | 70.88 | 135.35 | 49.01 | 88.89 | 0.5237 | 0.5514 |
| PHE | 187.59 | 248.85 | 38.66 | 87.37 | 0.7538 | 0.4425 |
| LEU | 159.14 | 219.49 | 37.32 | 88.26 | 0.7251 | 0.4229 |
| ILE | 159.82 | 220.11 | 35.32 | 87.57 | 0.7261 | 0.4034 |
| VAL | 133.94 | 194.60 | 36.68 | 87.27 | 0.6883 | 0.4202 |
| PRO | 130.02 | 192.85 | 39.65 | 91.42 | 0.6742 | 0.4337 |
| MET | 161.46 | 222.97 | 39.91 | 87.86 | 0.7241 | 0.4543 |
| TRP | 230.51 | 291.63 | 37.45 | 88.01 | 0.7904 | 0.4255 |
| GLY | 0.00 | 0.00 | 87.48 | 87.48 | 1.0000 | 1.0000 |
| SER | 85.27 | 148.71 | 45.90 | 87.71 | 0.5734 | 0.5234 |
| THR | 117.74 | 179.11 | 39.44 | 87.03 | 0.6574 | 0.4532 |
| TYR | 202.04 | 263.31 | 38.86 | 87.32 | 0.7673 | 0.4450 |
| GLN | 156.60 | 218.20 | 40.22 | 88.43 | 0.7177 | 0.4548 |
| ASN | 129.16 | 191.41 | 40.93 | 88.71 | 0.6748 | 0.4614 |
| ASP | 121.99 | 184.22 | 41.16 | 89.08 | 0.6622 | 0.4621 |
| GLU | 149.53 | 211.13 | 40.84 | 89.04 | 0.7082 | 0.4587 |
| HIS | 165.04 | 226.83 | 40.17 | 87.72 | 0.7276 | 0.4580 |
| LYS | 178.40 | 240.24 | 41.64 | 88.67 | 0.7426 | 0.4697 |
| ARG | 210.23 | 272.22 | 41.32 | 88.36 | 0.7723 | 0.4676 |
| CYS | 92.48 | 155.73 | 44.94 | 87.08 | 0.5939 | 0.5160 |

Table 1 shows that 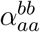 is significantly smaller than unity for all amino acids except glycine, as expected: the side chain shields the backbone from direct contact with the solvent, and the shielding is greater for bulkier side chains. These values are qualitatively similar to the 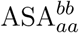 data of Creamer, and the fractional backbone exposure 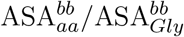 also coincides qualitatively with the ASA ratio used in the *Universal-backbone* model.

The shielding of the side chain by the backbone is also significant: 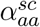 ranges from 0.524 for alanine to 0.790 for tryptophan, indicating that the backbone shields at least *∼*20% of even the bulkiest side chains. This effect was never explicit in previous models. Making it explicit is necessary here because the *Accessibility* model is constructed from the transfer free energy of the isolated side chain, in contrast to the other models, although this isolated side chain is implicitly incorporated in the definition of 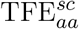 in eq 1. We compute the side-chain protection by the backbone explicitly, for consistency with the overall group definitions.

With these definitions, the transfer free energy of a protein is obtained, in the *Accessibility* model, by

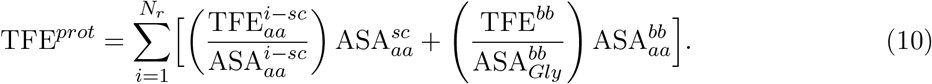

The second term of the sum is identical to that of the *Universal-backbone* model, and retains the chemically sound interpretation that all backbones contribute equally except for their degree of solvent exposure; the *Accessibility* model is therefore a “universal backbone” model. That TFE^*bb*^ refers to the isolated backbone needs no further specification, as this was uncontroversial in all models and derives directly from the experimental transfer free energies of glycine constructs.

The first, side-chain term does not match any of the current constructions, as both 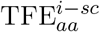 and 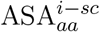 differ from the quantities in previous models. The *Universal-backbone* model can nevertheless be recovered qualitatively by multiplying both terms by 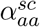, since 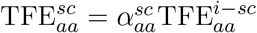 is the transfer free energy of the side chain shielded by the backbone. That transformation recovers the structure of the previous side-chain contributions, except that 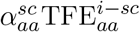 (eq 7) differs from what was considered to be 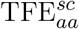 by the additional term 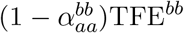. This term originates in the definition of 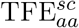 : subtracting TFE_*Gly*_ from TFE_*aa*_ in eq 1 removes a full backbone’s worth of free energy, while only the exposed part, considering side-chain shielding, should be removed. Multiplying TFE_*Gly*_ by 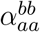 is not correct, because it would scale the end-group contributions. If TFE^*bb*^ is positive, as for protecting osmolytes, the additional term increases the total transfer free energy predictions; if negative, it decreases the side-chain contributions.

#### 2.2.2 Protecting osmolytes

Figure 3 shows the *Accessibility* model predictions of *m*-values for protecting osmolytes, compared to the available experimental data and to the *Established* model. Both models agree with experiment to the same extent, so the *Accessibility* model essentially reproduces the *Established* results. The same qualitative agreement holds for the denaturation of SH3 and dissociation of the GB1 dimer reported by Pielak and co-workers [24] (Supporting Information Section S5).

**Figure 3:**
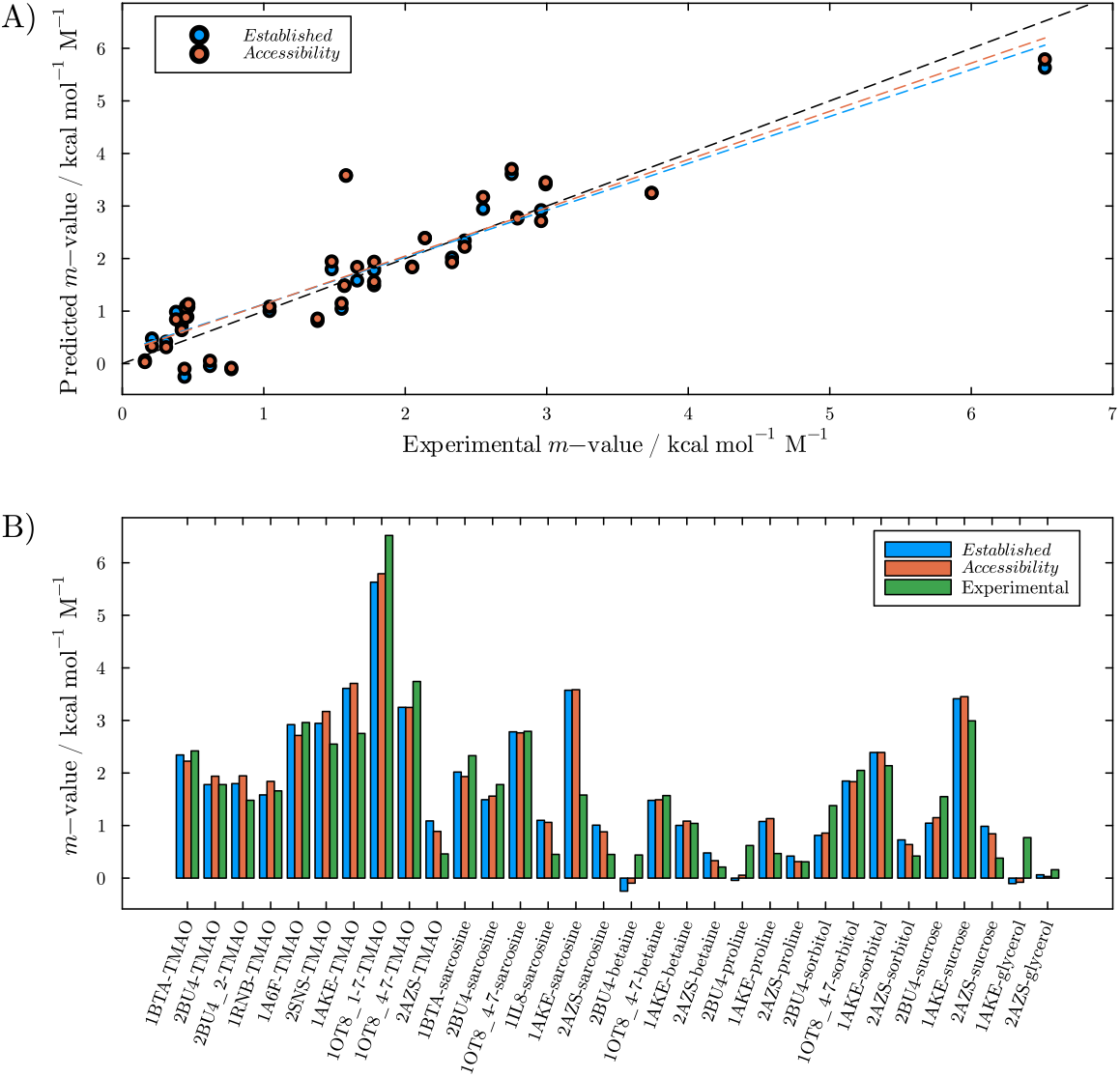
Experimental and predicted denaturation *m*-values for protecting osmolytes using the *Established* model and the *Accessibility* model. The predictions are similar and agree reasonably with the experimental data.

This agreement in the total *m*-value estimates indicates that both models capture the same average underlying amino acid contributions. Nevertheless, it occurs with a different balance of side-chain and backbone contributions, as shown in Figure 4 for TMAO (similar plots for other osmolytes are available as Supporting Information Figures S2–S10): the contributions of the side chains become relatively more important. For TMAO, backbone contributions are still greater than those of the side chains, but the side-chain contributions undergo a qualitative shift, becoming stabilizing in the *Accessibility* model. The *Accessibility* model thus provides the same (correct) predictions as the *Established* model for protecting osmolytes, with a new interpretation of backbone vs. side-chain contributions along the lines of the *Universal-backbone* model.

**Figure 4:**
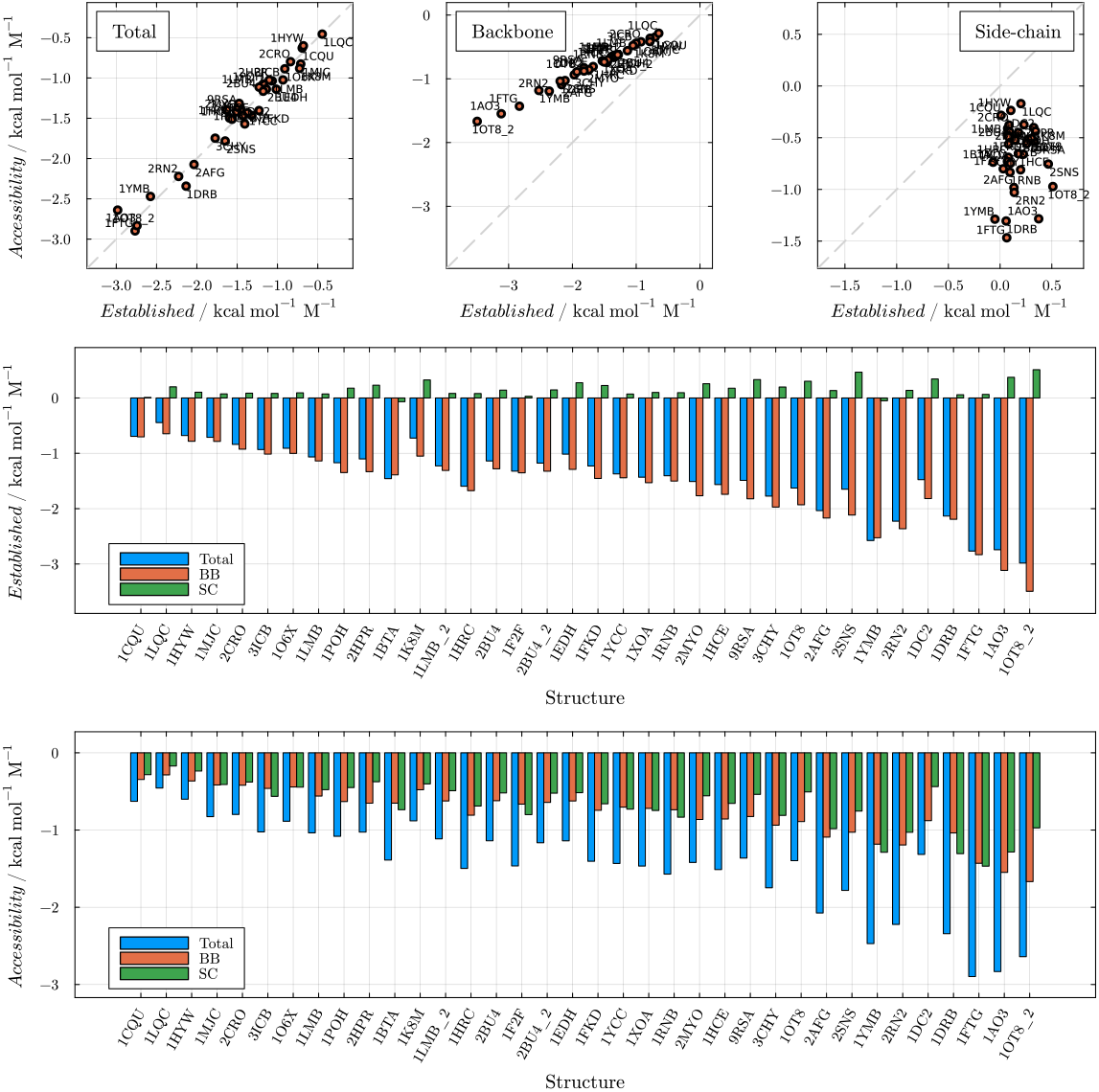
Total *m*-value predictions and backbone and side-chain contributions of the *Established* model and the *Accessibility* model for TMAO.

#### 2.2.3 Urea and the accessibility of the backbone

Figure 5A shows the *Accessibility* model predictions of *m*-values for urea-induced denaturation. The model overestimates the denaturing effect of urea, as the *Established* model did (Figure 1C), but the reason for the discrepancy is now of a different nature. As noted above, setting the accessibility parameter 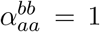 for all residues, instead of using the backbone exposure of each amino acid, reduces the model to the *Universal-backbone* model and recovers its predictive ability (Figure 5B).

**Figure 5:**
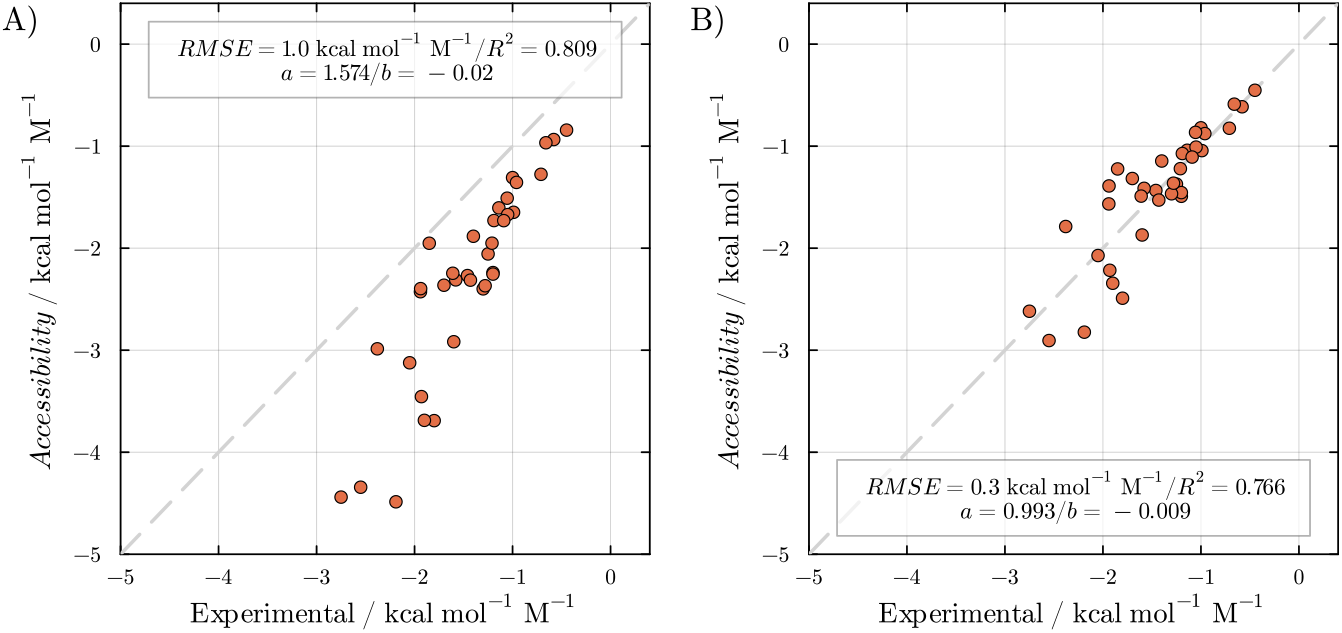
*Accessibility* model predictions of *m*-value of denaturation in urea, compared to experimental data using A) the ASA based 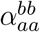 for each residue type and B) 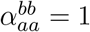 for all residues.

In this model, 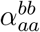 is the fraction of the backbone accessible surface area that is not shielded by the side chain. If the osmolyte is preferentially excluded and backbone–solvent interactions are non-specific, the accessible surface area estimates this accessibility. Urea, however, preferentially binds the protein and interacts with the backbone through hydrogen bonding, and since the carbonyl and amidic hydrogen of the backbone are on the opposite side of the side chain, the surface area may not be a good proxy for the side-chain shielding.

Figure 6 shows the number of hydrogen bonds of the backbone of different amino acid residues in 1 mol L*−*1 aqueous urea, relative to that of glycine. The backbones of most residues form a similar number of hydrogen bonds with urea as glycine, meaning that backbone hydrogen bonding is not effectively shielded by the side chains, and that the surface area ratios overestimate this shielding. These results are similar to those of Best and co-workers in a simulation of an intrinsically disordered protein [19]. Accounting for this by increasing the backbone accessibility relative to the surface area ratios, the model captures the experimental denaturation *m*-values. Using the exact hydrogen-bond ratio of Figure 6 as *α*^*bb*^ for each residue type, we obtain *R*^2^ = 0.772 (Supplementary Information Figure S17), similar to unitary accessibility for all residues, which is thus a reasonable approximation. The *Universal-backbone* model is therefore a particular case of the *Accessibility* model in which the specific and directional nature of the preferential binder–backbone interactions is taken into account.

**Figure 6:**
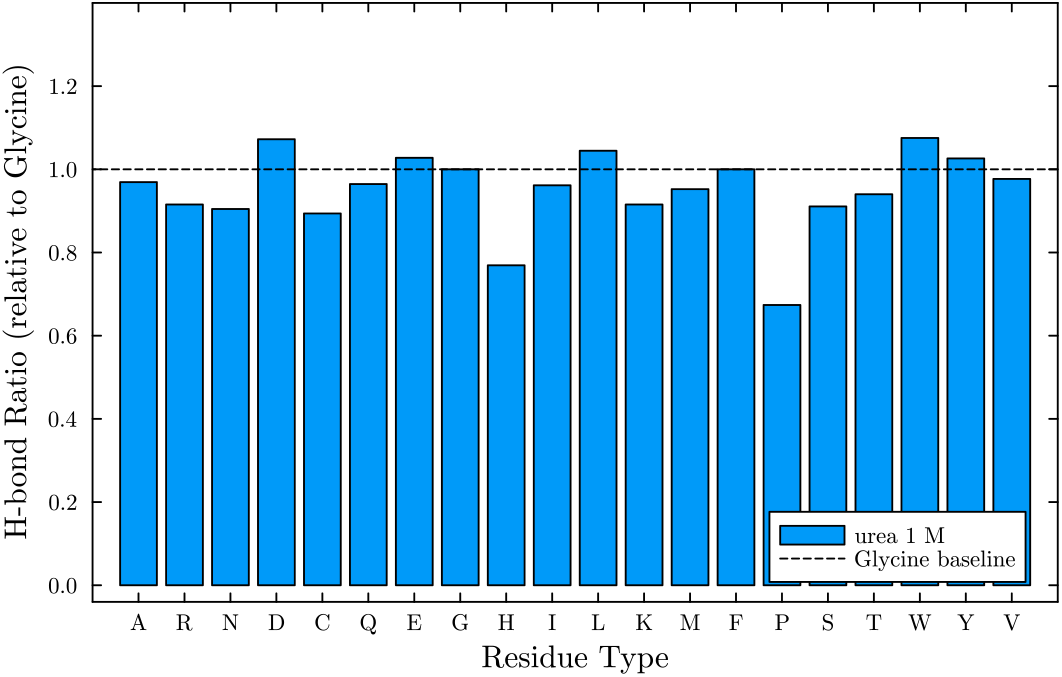
Number of hydrogen bonds of urea with the backbone of different amino acid types, relative to Glycine, in simulations of free Ace/NH2 capped peptides (results for non-capped peptides, either charged or neutral are similar and available in Supporting Information Section S9).

We also tried to obtain optimal *α*^*bb*^ parameters by fitting the model to the 36-protein model set of Figure 5. To avoid excessive overfitting, we adjusted only the parameters of the bulkier hydrophobic side chains, those of phenylalanine, tyrosine, and tryptophan. Only slightly better fits are obtained relative to *α*^*bb*^ = 1, and with unphysical compensation of accessibilities (*R*^2^ = 0.79 with 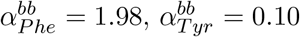, and 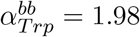). Restricting *α*^*bb*^ *<* 1.2, the best fit (*R*^2^ = 0.77) is obtained with 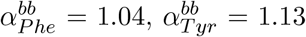, and 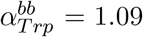), supporting the full-accessibility model. In fact, if non-specific interactions with side chains cooperate with backbone hydrogen bonding, *α*^*bb*^ can exceed one, as for some residues in Figure 6. Of course, some protecting cosolvents can form hydrogen bonds (sugars and polyols), but they are known to be preferentially excluded from the protein surface such that the water accessibility can be used as a proxy for the mutual shielding.

## 3. Microscopic Validation With Molecular Simulations

Predictions of denaturation *m*-values rely on assumptions about the structure of the denatured states, which are ultimately empirical from agreement of denaturation *m*-values with experimental data. On the other hand, molecular dynamics simulations and the Kirkwood-Buff theory of solutions provide a framework to compute differences in transfer free energies between any two structures, given adequate force-fields and proper sampling [16, 19, 25, 26, 27]. Simulations have validated the decomposition of backbone and side-chain roles for urea [18, 16, 19, 25], agreeing with the balanced contributions of the *Universal-backbone* model; the *Accessibility* model provides the same predictions in this case. The decomposition of the stabilizing effects of protectants, nevertheless, has not been investigated with simulations. Here we evaluate the three models against molecular dynamics simulations of protein denaturation and protein-protein association in aqueous TMAO, assigning solvent density fluctuations to the closest protein atom, in a Voronoi spatial decomposition scheme, as proposed in [19].

First, a set of denatured models were simulated to evaluate the total predictions and backbone and side-chain contributions associated with variable exposed surface areas; second, a set of dimers for which the *total m*-values of dissociation predicted by the models differ was considered. This set is crucial because, as discussed, the models can predict similar total transfer free energies despite different backbone and side-chain weights. By studying transformations with different total predictions, the models can be compared before invoking any group or spatial decomposition scheme. Agreement of the total predictions with the simulations validates the model construction to the extent that the force-fields are accurate, and also provides a set of systems for which experimental validation is in principle possible.

Figures 7A-D show the total, backbone, and side-chain contributions obtained with each additive model vs. the MD predictions for the denaturation of the B-domain of protein A. Each dot in Figures 7A-C represents a different denatured state obtained with structure-based models and simulated atomistically in 0.5 mol L^*−*1^ TMAO (simulation details are given in a previous publication [13]). Larger *m*-values correspond to models with greater denaturation extent relative to the native structure.

**Figure 7:**
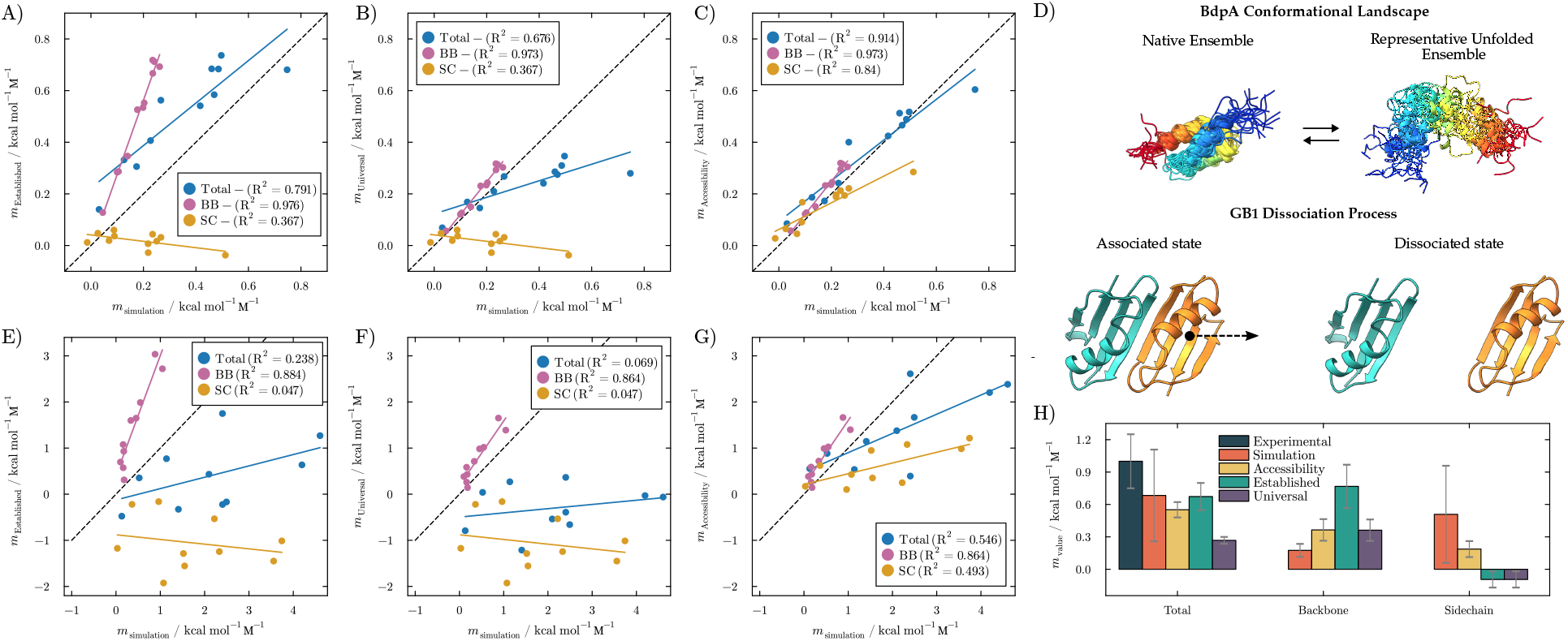
Model predictions and molecular simulations estimates for denaturation and dimer dissociation in TMAO. A-D) *m*-values of BdpA unfolding for various denatured ensembles. E-G) Variation in transfer free energies induced by TMAO, for the dissociation of oligomers. H) *m*-values of dimer dissociation in TMAO for the GB1 dimer. Only the *Accessibility* model displays qualitative agreement with the decomposition of backbone and side-chain contributions to transfer free energies.

The *Established* model slightly overestimates the total *m*-values, the *Universal-backbone* model underestimates them, particularly for the most denatured states, and the *Accessibility* model gives the most coincident totals. While this appears to favor the new model, it depends on protein composition and is not the most relevant aspect of the comparison. More importantly, the relative contributions of backbones and side chains differ substantially from the simulation decomposition: the *Established* model strongly overestimates the backbone contributions and underestimates the side-chain ones, and while the *Universal-backbone* model fixes the backbone contributions, it fails to represent the weight of side chains. Both backbone and side-chain contributions of the *Accessibility* model, in contrast, agree well with the simulations. The simulations therefore support the *Accessibility* model construction for the roles of backbones and side chains in protein denaturation, independently of the denatured-state model, which is a free parameter in the direct *m*-value computation from the native structure.

Figures 7E-H show the model vs. simulation predictions for dissociation transfer free energies in TMAO, for representative dimeric associations where the model predictions differ substantially. These are biophysically characterized homodimers with different interface architectures (Supplementary Information Section S8) [30, 31, 32, 33, 34, 35, 36, 37, 38, 39, 24] whose interfaces provide structurally distinct tests of the predicted backbone and side-chain contributions upon dissociation. In most cases (except GB1-A34F), the interfaces are dominated by helical motifs or side-chain-rich contacts. Thus, dissociation exposes predominantly side-chain surfaces and leads to larger *m*-values in the *Accessibility* than in the *Established* model, reaching differences of approximately 1–2 kcal mol^*−*1^ M^*−*1^ for TMAO in the most divergent cases, with similar differences for other osmolytes. The GB1-A34F dimer, in contrast, contains paired *β*-strands at the interface, increasing the backbone contribution and leading to a larger prediction in the *Established* than in the *Accessibility* model.

As for protein denaturation, the *Universal-backbone* model clearly improves the characterization of backbone contributions relative to the *Established* model, but the role of side chains is qualitatively incorrect. The *Accessibility* model qualitatively fixes the side-chain contributions, which become positive (favoring dimer stability). Still, with the current simulation model and decomposition strategy, the simulations predict even more positive side-chain contributions than the *Accessibility* model, and the total *m*-values of all models are lower than the simulated ones. Figure 7H illustrates these differences for the GB1-A34F dimer, for which the total experimental dissociation transfer free energy is available [24]. The simulation, *Accessibility*, and *Established* models all somewhat underestimate the experimental value, although within the observed uncertainties. As a representative case, the total simulated TFE results from positive backbone and side-chain contributions, qualitatively consistent with the *Accessibility* model. The *Established* model obtains a similar total TFE but from the compensation of an overestimated backbone contribution and a negative side-chain contribution, while the *Universal-backbone* model significantly underestimates the total TFE because of its underestimated side-chain contribution. Quantitative differences between the *Accessibility* model and the simulations for the side-chain contributions may reflect inaccuracies of the force-fields, or of the experimental transfer free energies of specific amino acid types, and have greater impact in the total *m*-values of dimer dissociation.

One limitation of this analysis is that the 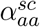 and 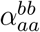 values in Table 1 are averages over the entire CATH S20 database, and therefore are insensitive to specific conformation of each residue. The actual solvent exposure of a given residue depends on its local structural context, as demonstrated explicitly by Thirumalai and co-workers [58, 59]. This limitation is shared by all additive models. Notably, our own atomistic simulations (Figure 7), which do resolve conformation-specific solvent structure through the Kirkwood–Buff decomposition, independently support the backbone/side-chain balance predicted by the *Accessibility* model across a structurally diverse ensemble of denatured states and dimer interfaces — suggesting that, in aggregate, the database-averaged accessibility parameters capture the dominant physical effect despite neglecting residue-level fluctuations.

These and other oligomeric structures with notable model discrepancies could be used to validate experimentally, and in a definitive manner, this or other proposals on the distribution of backbone vs. side-chain contributions. This structural sensitivity highlights a major utility of the *Accessibility* model: because it correctly partitions the free energy weights, it can serve as a diagnostic tool to predict how specific mutations at oligomeric interfaces alter osmolyte-induced shifts in assembly equilibria. For engineering protein therapeutics or understanding the cellular preservation of macro-molecular complexes under osmotic stress, shifting from a generic “backbone protection” view to a residue-specific “cooperative surface exclusion” view opens new avenues for rational protein design.

## 4. Osmophobic Side Chains Are Fundamental for Protecting Osmolytes

Having reconciled the total *m*-value predictions of the *Established* and *Universal-backbone* models with the experimental data, we can reevaluate the distribution of backbone and side-chain contributions to cosolvent-induced effects on protein folding. For urea, a strongly denaturing osmolyte, the interpretation follows that of Moeser and Horinek and the *Universal-backbone* model: side chains and backbone contribute similarly to the overall denaturing effect, with urea displaying favorable interactions with both groups that lead to preferential solvation.

The balanced redistribution of effects is now extended to protecting osmolytes, challenging the established osmophobic interpretation [3], as anticipated in Figure 4 for TMAO. Averaging the effects per residue over the database of protein models studied yields general rules: Table 2 shows that, for strong protecting osmolytes, the contributions per residue to *m*-values are of the same order for backbones and side chains. These interactions are unfavorable, so osmophobicity and preferential exclusion can be attributed to both groups, with the backbone or the side chain being quantitatively more important depending on the osmolyte (and on the residue composition). Osmophobicity is thus a cooperative effect of side chains and backbone.

**Table 2.** Mean denaturation *m*-values per residue, in cal mol^*−*1^ M^*−*1^, for different osmolytes, and backbone and side-chain contributions. Values are averaged over the CATH S20 protein set; uncertainties (*±*) are standard deviations across proteins.

| Osmolyte | Total | Backbone | Side chains | Effect |
| --- | --- | --- | --- | --- |
| trehalose | $27.22 \pm 4.83$ | $10.88 \pm 1.37$ | $16.34 \pm 4.40$ | strong protectant |
| TMAO | $22.48 \pm 3.41$ | $15.79 \pm 1.99$ | $6.69 \pm 2.25$ | strong protectant |
| sarcosine | $19.55 \pm 2.72$ | $9.12 \pm 1.15$ | $10.43 \pm 2.15$ | strong protectant |
| sucrose | $18.90 \pm 3.95$ | $10.88 \pm 1.37$ | $8.02 \pm 3.56$ | strong protectant |
| sorbitol | $13.06 \pm 2.38$ | $6.14 \pm 0.78$ | $6.92 \pm 2.07$ | strong protectant |
| proline | $6.18 \pm 2.97$ | $8.42 \pm 1.06$ | $-2.24 \pm 2.86$ | medium protectant |
| betaine | $5.29 \pm 4.53$ | $11.76 \pm 1.48$ | $-6.47 \pm 4.43$ | weak protectant |
| glycerol | $-0.36 \pm 2.14$ | $2.46 \pm 0.31$ | $-2.81 \pm 2.12$ | mixed |
| urea | $-12.73 \pm 2.05$ | $-6.84 \pm 0.86$ | $-5.89 \pm 1.50$ | strong denaturant |

For osmolytes with mixed effects (qualitatively dependent on protein composition), the small absolute *m*-values result from opposite compensating effects, with exclusion at the backbone opposing favorable osmolyte–side-chain interactions, consistent with previous simulations, as for betaine [29]. These osmolytes thus follow the conventional backbone-centered osmophobic interpretation. Finally, for urea, cooperative favorable interactions occur at both groups.

## 5. Conclusions

Here we show that an impasse in additive transfer models arises from an incomplete treatment of the mutual shielding between backbone and side-chain groups when extracting group transfer free energies from experimental data. Introducing a new *Accessibility* model, we account for both the shielding of the backbone by the side chain and the shielding of the side chain by the backbone. For protecting osmolytes, the model reproduces the predictive accuracy of the *Established* model while providing a fundamentally different — and physically grounded — decomposition of backbone and side-chain contributions. The model reveals that the ASA-based backbone accessibility does not account for the preferential binding and hydrogen-bonding nature of urea–backbone interactions. This mechanism-dependent interpretation of backbone accessibility — geometric for excluded cosolvents, complete for preferential binders — unifies the treatment of denaturants and protectants in a single framework.

The revised decomposition reinterprets the osmophobic effect. Rather than being driven exclusively by unfavorable backbone–osmolyte interactions, protein stabilization by strong protecting osmolytes involves cooperative osmophobic contributions from both backbone and side chains. For TMAO, sarcosine, sucrose, trehalose, and sorbitol, side-chain contributions are comparable to or exceed those of the backbone. This extends osmophobicity to the full protein surface and challenges the view that the backbone is the sole mediator of osmolyte-induced stabilization.

A three-tier classification of osmolyte action emerges. Strong protectants exhibit cooperative backbone and side-chain stabilization. Weak or borderline osmolytes, such as proline, betaine, and glycerol, display competing contributions — backbone exclusion opposed by favorable side-chain interactions — giving small net effects sensitive to protein composition. Denaturants such as urea show cooperative destabilization by both groups. This classification provides a mechanistic basis for why certain osmolytes are more effective stabilizers than others, and suggests that stabilization strategies should consider the amino acid composition of the exposed surface, not only the extent of backbone exposure.

## Materials and Methods

### Transfer free energy data

Apparent side-chain transfer free energies 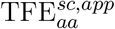 and backbone transfer free energies TFE^*bb*^ for nine cosolvents (TMAO, sarcosine, betaine, proline, sorbitol, sucrose, urea, glycerol, and trehalose) were taken from Auton and Bolen [12, 15]. The Glycine activity correction for urea, *γ*_*Gly*_ = *−*14.47 cal mol^*−*1^ M^*−*1^, was taken from Moeser and Horinek [16]. For all other cosolvents the Glycine activity corrections are expected to be negligible and were set to zero.

Accessible surface areas (ASAs) were computed using a Fibonacci-lattice dot-counting algorithm equivalent to the Shrake–Rupley method [40], as implemented in the `PDBTools.jl` package [41]. A probe radius of 1.4 Å and 512 surface points per atom were used throughout. Atomic radii were assigned by hybridization type following the united-atom parameterization of Creamer et al. [14] for all but *Record* models. Neighbor lists required for the pairwise exclusion step were computed with the `CellListMap.jl` package [42]. For *Record* models, the ASAs were computed with Richard’s radii and taking into account occluded cavities [43], implemented in PDBTools.jl v3.39.0 to match previous results [21].

The accessibility parameters 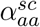 and 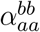 (Table 1) were computed from the CATH S20 non-redundant protein domain database (release downloaded May 2026) [44]. For every occurrence of each amino acid type, four ASA values were computed: the side-chain ASA in the intact residue 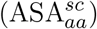, the side-chain ASA with all backbone atoms removed 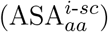, the backbone ASA in the intact residue (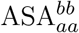, in the presence of the backbones of immediately neighboring residues), and the backbone ASA with all side-chain atoms removed (ASA^*bb*^). The accessibility fractions are the averages of these ratios over the database. Because they reflect the intrinsic local geometry between the side chain and backbone of each residue type — rather than the global fold — the values apply to both native and denatured states and are largely insensitive to the non-redundancy threshold; comparable values are obtained with the earlier Creamer dataset [14].

Crystal structures for all benchmark proteins were retrieved from the Protein Data Bank (PDB) [45]. Hydrogen atoms, solvent molecules, and non-protein chains were removed before any calcu-lation, using `PDBTools.jl` [41]. The change in ASA upon unfolding, ΔASA, was estimated with the mean denatured-state model of Creamer et al. [14] The denatured-state ASA was cross-checked against the web server at best.bio.jhu.edu/mvalue, with equivalent predictions; results reported here use the Creamer model (server output type 2, corresponding to average denatured-state ASAs), following established procedures [12, 15].

For urea, the benchmark set comprised proteins with experimental denaturation *m*-values published in refs. [12, 16]: 1CQU, 1LQC, 1HYW, 1MJC, 2CRO, 3ICB, 1O6X, 1LMB, 1POH, 2HPR, 1BTA, 1K8M, 1HRC, 2BU4, 1F2F, 1EDH, 1FKD, 1YCC, 1XOA, 1RNB, 2MYO, 1HCE, 9RSA, 3CHY, 1OT8, 2AFG, 2SNS, 1YMB, 2RN2, 1DC2, 1DRB, 1FTG, and 1AO3 (PDB IDs). For protecting osmolytes (TMAO, sarcosine, betaine, proline, sorbitol, sucrose, trehalose, and glycerol), a subset of these structures with available experimental data was used as compiled in ref. [15], including PDB entries 1BTA, 2BU4, 1RNB, 2SNS, 1AKE, 1OT8, and 1IL8. Further details are available in the Supplementary Information.

### Molecular Dynamics Simulations

The protein-water-TMAO systems were modeled using the CHARMM36 [46] force field for the proteins and the TIP3P model [47] for water. TMAO was described using the Netz force field [48] with the Shea group’s scaling correction [49] to ensure the accurate reproduction of Kirkwood–Buff integrals (KBIs). The initial configurations were constructed with Packmol [50, 51], and simulations were performed in GROMACS 2021.2 [52] in the NPT ensemble at 298.15 K and 1 atm.

BdpA (PDB id. 1BDD) [53] denaturation was studied by first simulating a structure-based model at the folding temperature [54]. Each of 5000 models was reconstructed atomistically and solvated in explicit solvent. Then, MD simulations were performed for the characterization of the solvent structure and thermodynamics through Kirkwood-Buff theory (see Supplementary Information Section S7). Minimum-distance counts were used to compute distribution functions and KB integrals [28, 55]. The bulk-solvent non-ideality for TMAO was obtained by fitting the concentration-dependence of *m*-values reported previously [56, 13]. The structures were clustered by distance-similarity [57], and for each cluster the average *m*-values were obtained, and are represented as a data point in Fig. 7. All details of the simulations and solvation analyses are described in previous publications [56, 13]. Dimer dissociation *m*-values were computed by simulating the assembled dimer and the monomers independently in 1 mol L^-1^ solutions of TMAO. Further details can be found in Supplementary Information sections S6 and S7.

Kirkwood-Buff integrals (KBIs) were computed using the ComplexMixtures.jl package [55], and the decomposition of the KBIs into backbone and side-chain contributions was obtained in the newly implemented `type=:kbi` option of the `contributions` function in ComplexMixtures.jl version 2.18.0.

## Data availability and reproducibility

All transfer models (Auton–Bolen (*Established*), Moeser–Horinek (*Universal-backbone*), Record, and *Accessibility*) and ASA calculations are fully implemented in `PDBTools.jl` [41] (v3.39.0 or later) such that calculations can be directly reproduced without any third-party application. The structures of dimers are available in the protein data bank, and the denatured models are available in the Zenodo repository https://doi.org/10.5281/zenodo.17155226 [13]. Full MD trajectories with cosolvents are available upon request to authors.

## Acknowledgments

The authors acknowledge the financial support of FAPESP (2018/24293-0, 2013/08293-7, 2019/17874-0, 2020/04549-0, 2023/13899-3, 2025/16377-3, 2025/03933-5), and CNPq (301909/2022-9). Research developed with the help of CENAPAD-SP (National Center for High-Performance Processing in São Paulo), project UNICAMP/FINEP–MCTI, and the Coaraci Supercomputer.

## Author contributions

A.F.P. performed MD simulations of denatured models, H-bond occupancy, analyzed data, and wrote the paper. J.O.A. performed the simulations of dimers, W.T. selected models for analyses and computed predictions, and L.M. designed research, proposed and implemented the Accessibility model, analyzed data, and wrote the paper.

## Competing interests

The authors declare no competing interest.

## Supporting Information

The Supporting Information contains supplementary methods and results supporting the main text: comparison of the *Universal-backbone* and *Established* models for total, backbone, and side-chain *m*-value predictions in urea (Fig. S1, Section S1); comparisons of the *Accessibility* and *Established* models for total, backbone, and side-chain *m*-value predictions across all nine cosolvents studied (Figs. S2–S10, Section S2); comparisons of the *Universal-backbone* vs. Record models for urea (Fig. S11, Section S3) and of the Record and *Established* models for protecting osmolytes (Fig. S12, Section S4); validation of the *Accessibility, Established, Universal-backbone*, and Record models against experimental denaturation and dimer-dissociation free energies for SH3 and GB1 (Figs. S13 and S14, Section S5); comparison of Record model predictions with molecular dynamics simulations of TMAO-induced folding and dimer dissociation (Fig. S15, Section S6); molecular dynamics simulation protocols (Section S7); structural details of the protein dimers used for *m*-value comparisons (Section S8); and the analysis of backbone hydrogen-bond shielding by side chains in urea (Figs. S16 and S17, Section S9). SI References are also provided.

## Notes

### Competing Interest Statement

The authors have declared no competing interest.

### Summary of Updates

Updated simulations with new dimers, add comparison to Record models, revised text.

