## Supplementary Information for "Extending the osmophobic effect to protein side chains with a physical transfer model across osmolyte classes"

S1. Predictions of *m*-values, and backbone and side-chain contributions, of the Universal-backbone model vs. the Established model for urea. 2

S2. Predictions of *m*-values, and backbone and side-chain contributions, of the Accessibility model vs. the Established model for all osmolytes. 3

S3. Predictions of *m*-values, and backbone and side-chain contributions, of the Universal-backbone vs Record models for urea. 12

S4. Predictions of *m*-values of the Established and Record models for protecting osmolytes. 13

S5. Transfer free energies of denaturation and dimer dissociation for SH3 and GB1, using the Accessibility, Established, Universal-backbone and Record models. 14

S6. Record model predictions for TMAO in comparison with molecular dynamics simulations 17

S7. Molecular Dynamics Simulations Protocols 19

S8. Model predictions in selected dimers 21

S9. Shielding of urea-backbone hydrogen bonds by side chains 22

References 25

### S1. Predictions of *m*-values, and backbone and side-chain contributions, of the *Universal-backbone* model vs. the *Established* model for urea.

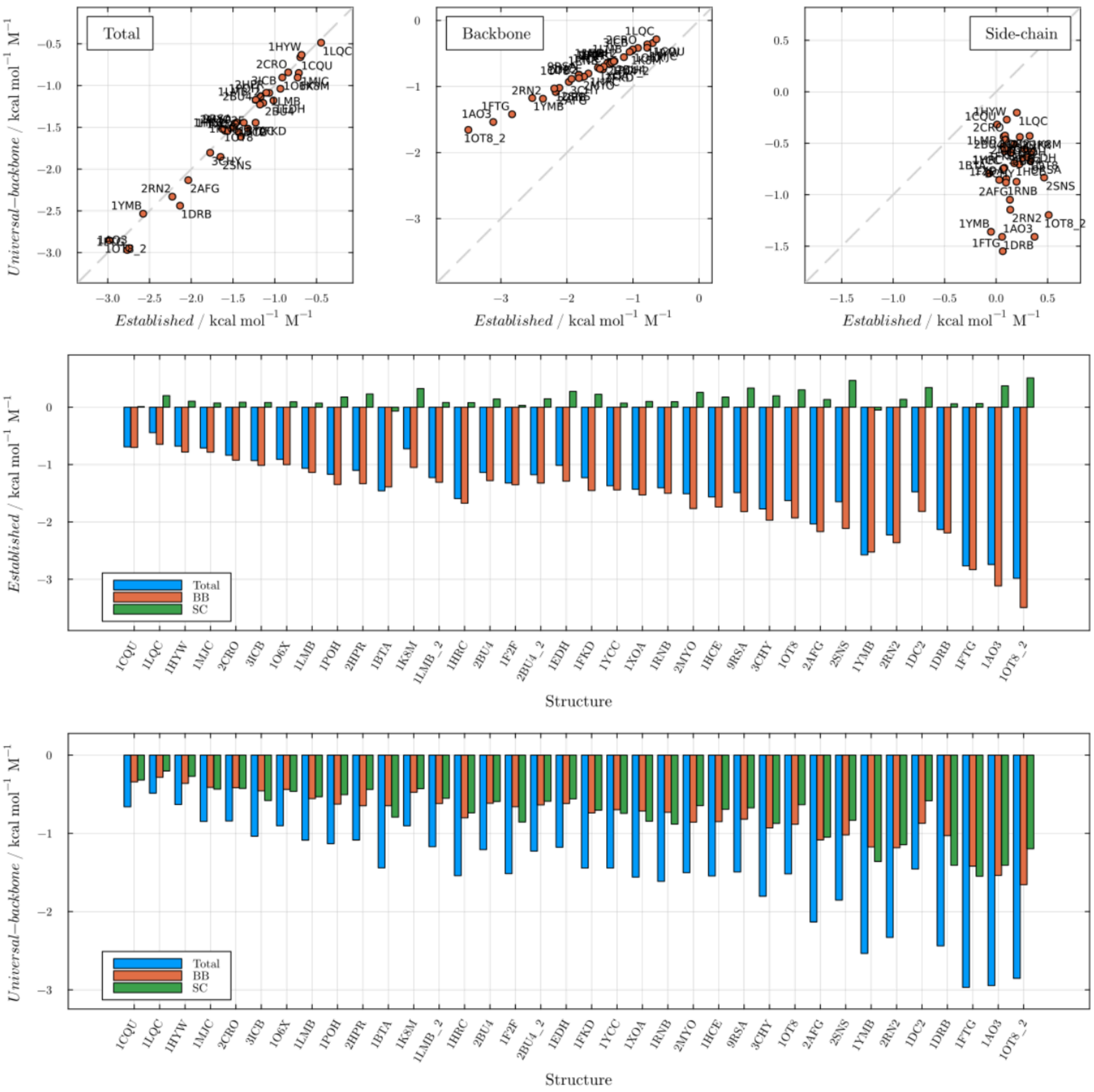


**Figure S1.** Predictions of the *Universal-backbone* and *Established* models for total, backbone and side-chain *m*-values in urea.

### S2. Predictions of *m*-values, and backbone and side-chain contributions, of the *Accessibility* model vs. the *Established* model for all osmolytes.

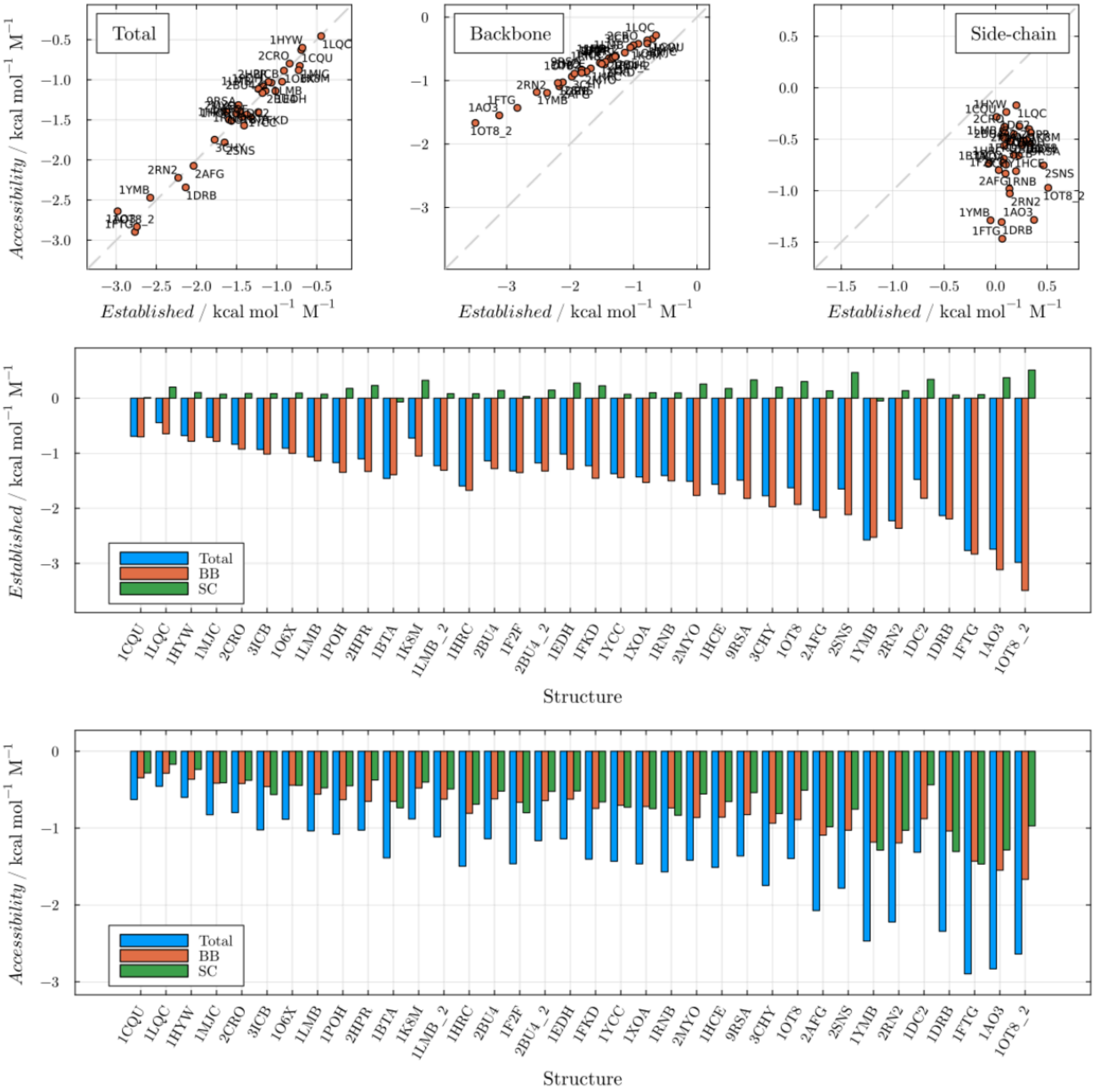


**Figure S2.** Predictions of the *Established* and *Accessibility* models for total, backbone and side-chain *m*-values in urea.

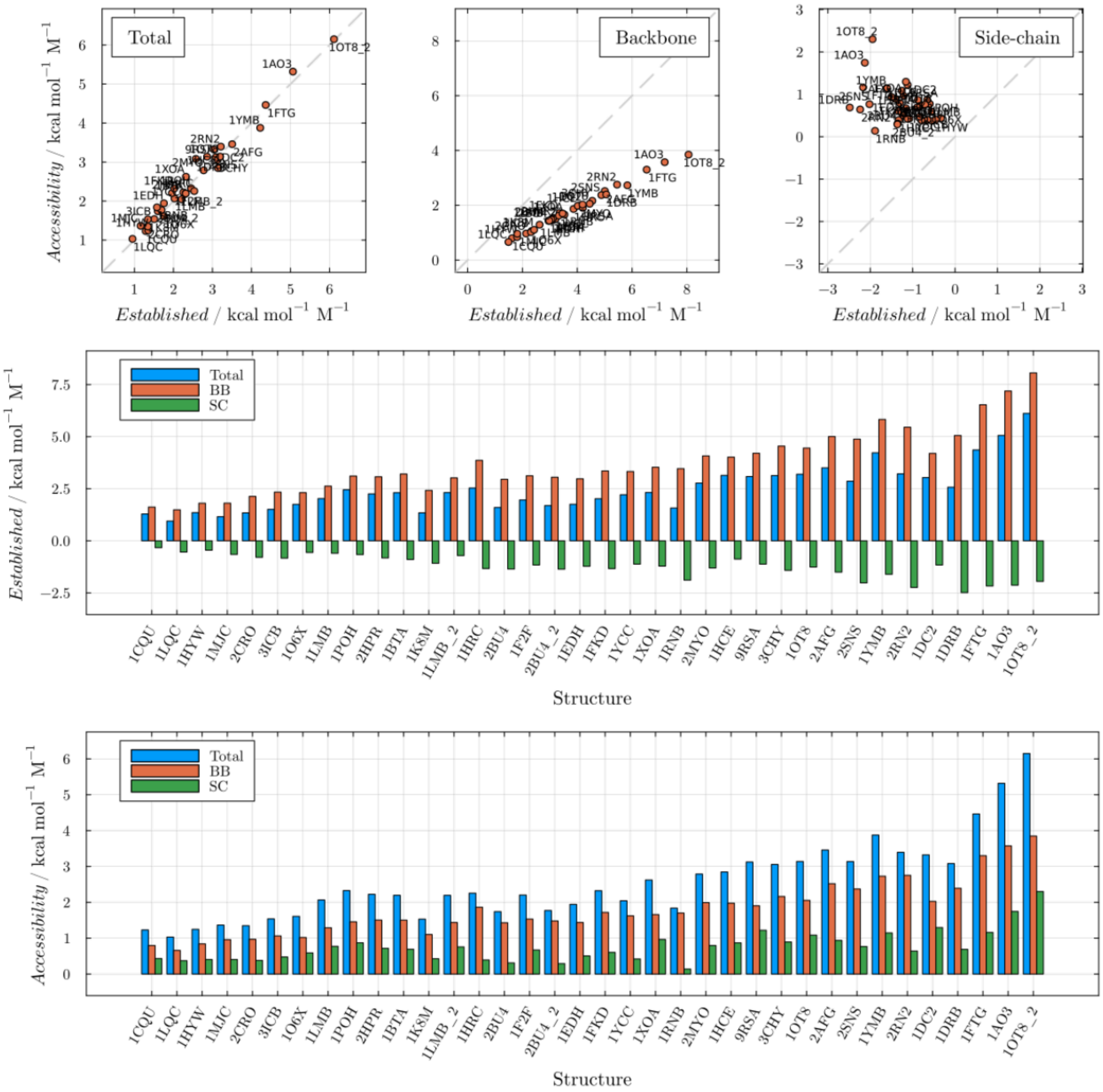


**Figure S3.** Predictions of the Es*tablished* and *Accessibility* models for total, backbone and side-chain *m*-values in TMAO.

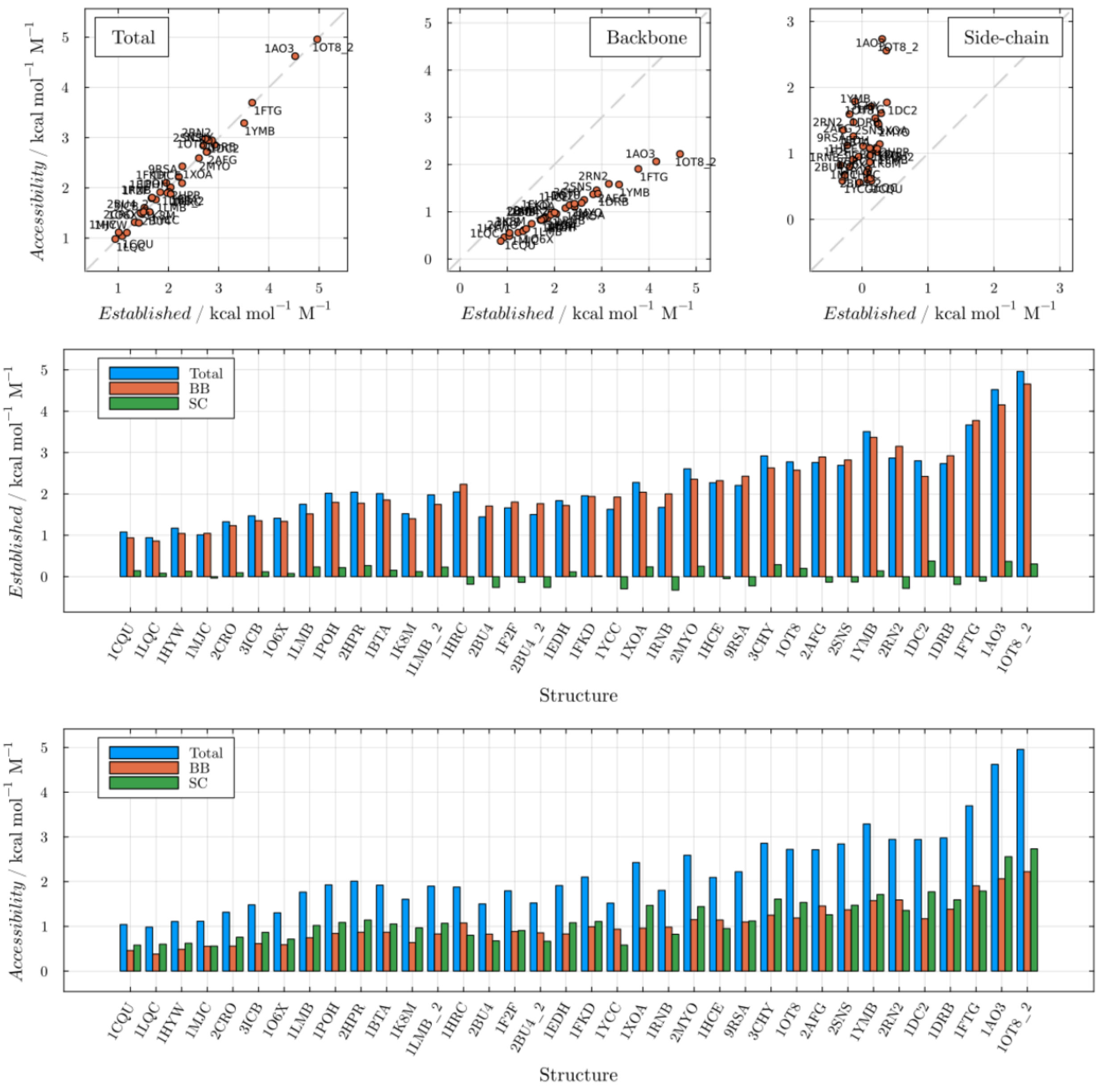


**Figure S4.** Predictions of the *Established* and *Accessibility* models for total, backbone and side-chain *m*-values in sarcosine.

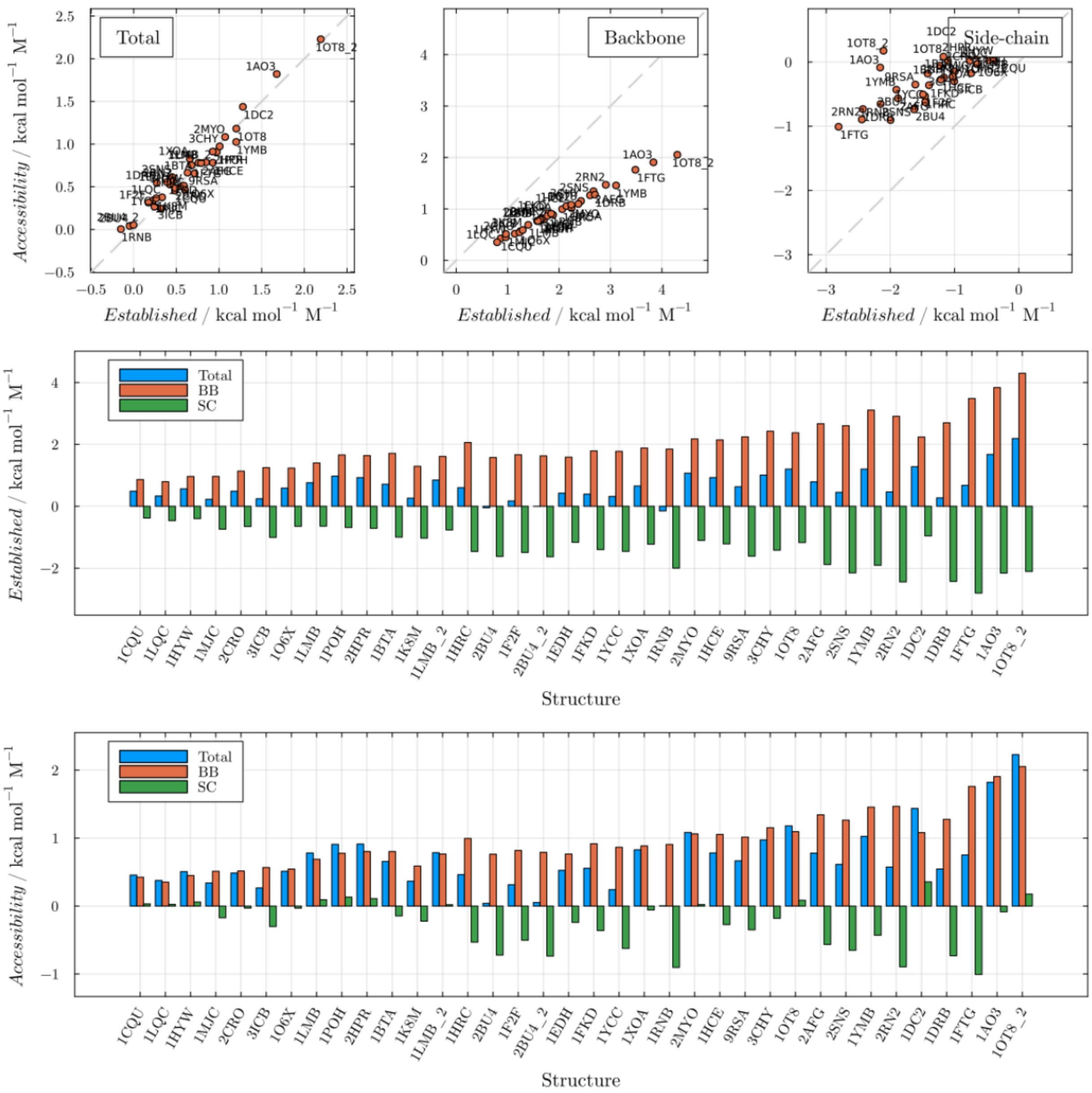


**Figure S5.** Predictions of the *Established* and *Accessibility* models for total, backbone and side-chain *m*-values in proline.

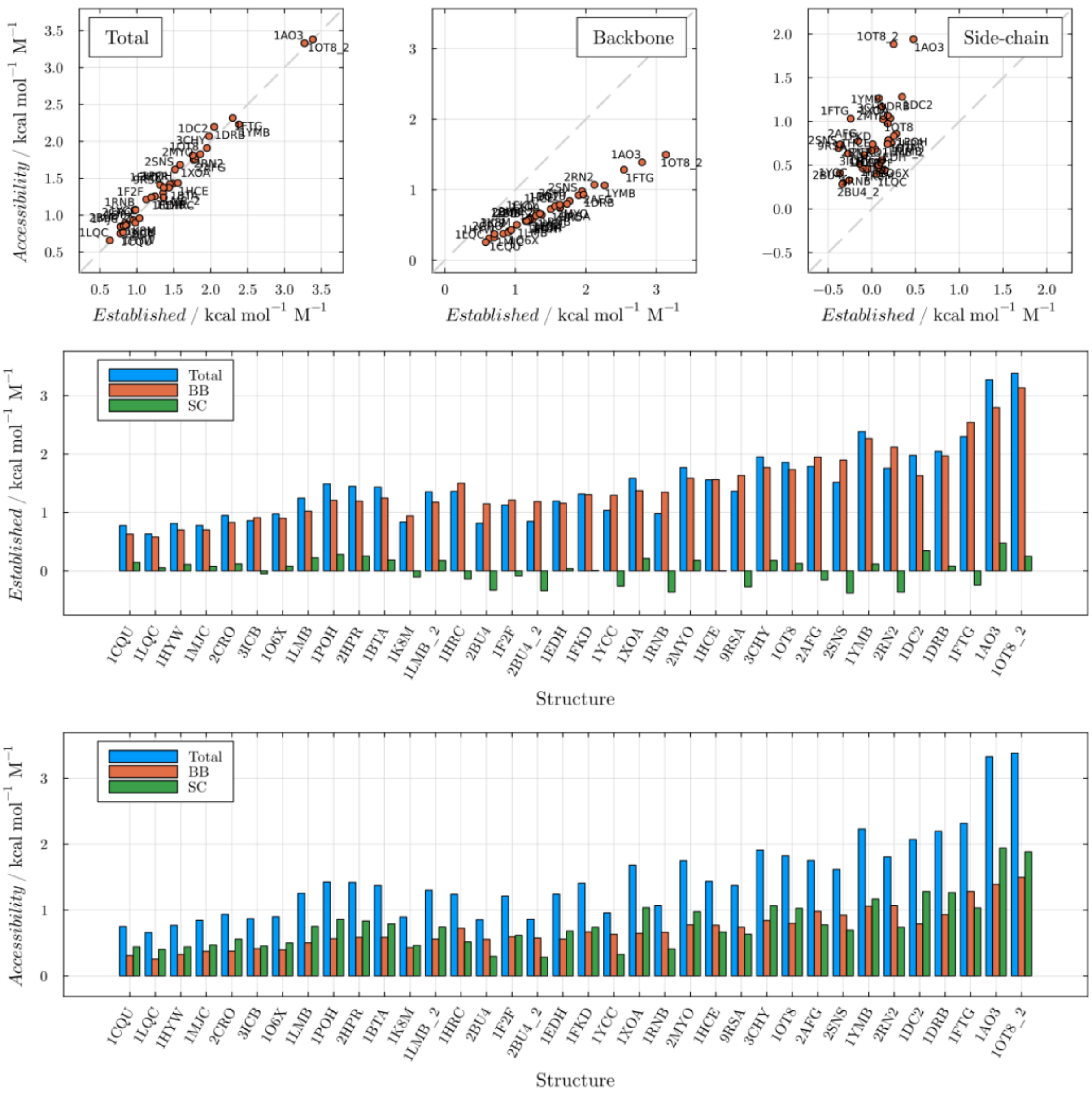


**Figure S6.** Predictions of the *Established* and *Accessibility* models for total, backbone and side-chain $m$-values in sorbitol.

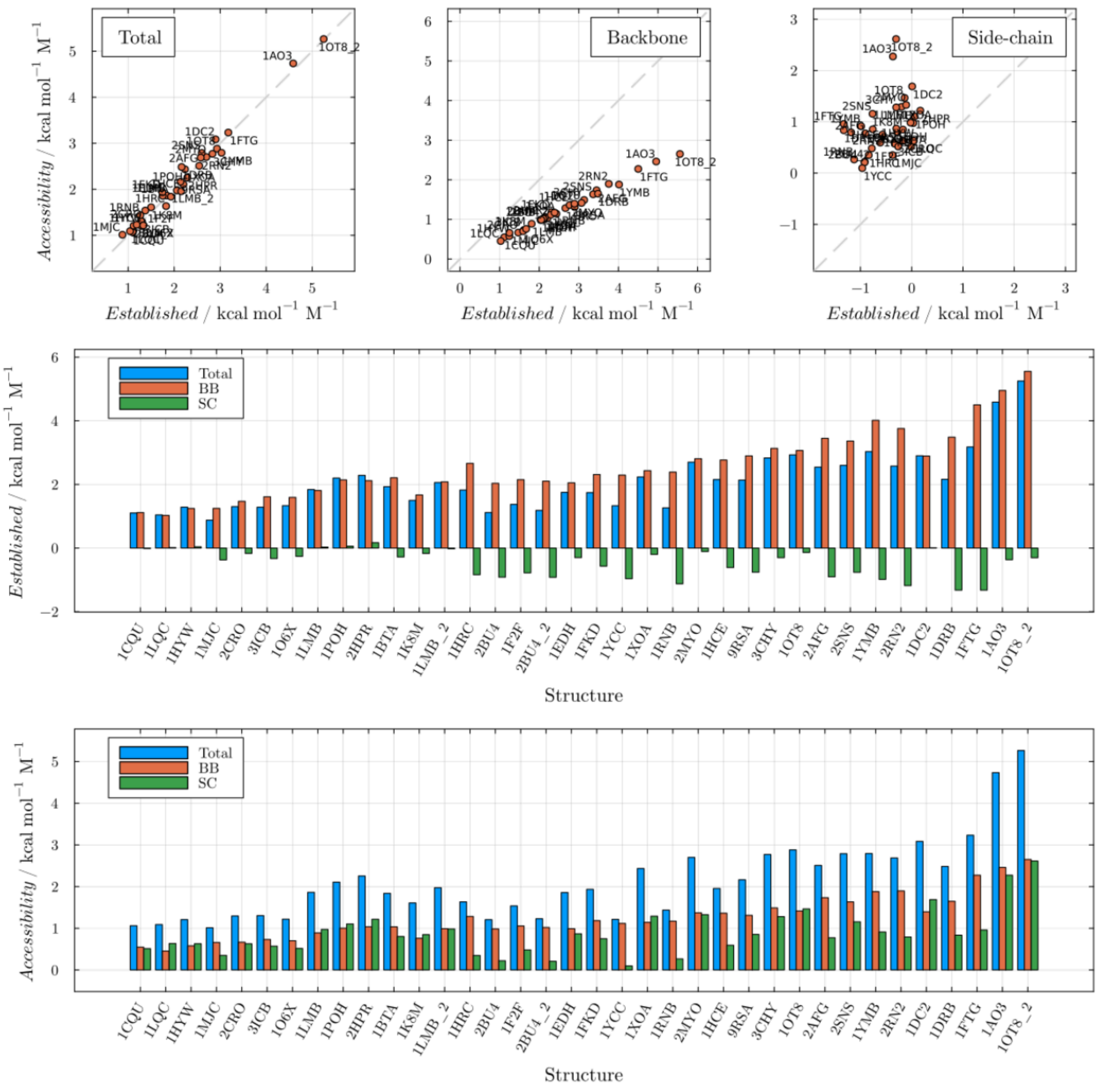


**Figure S7.** Predictions of the *Established* and *Accessibility* models for total, backbone and side-chain *m*-values in sucrose.

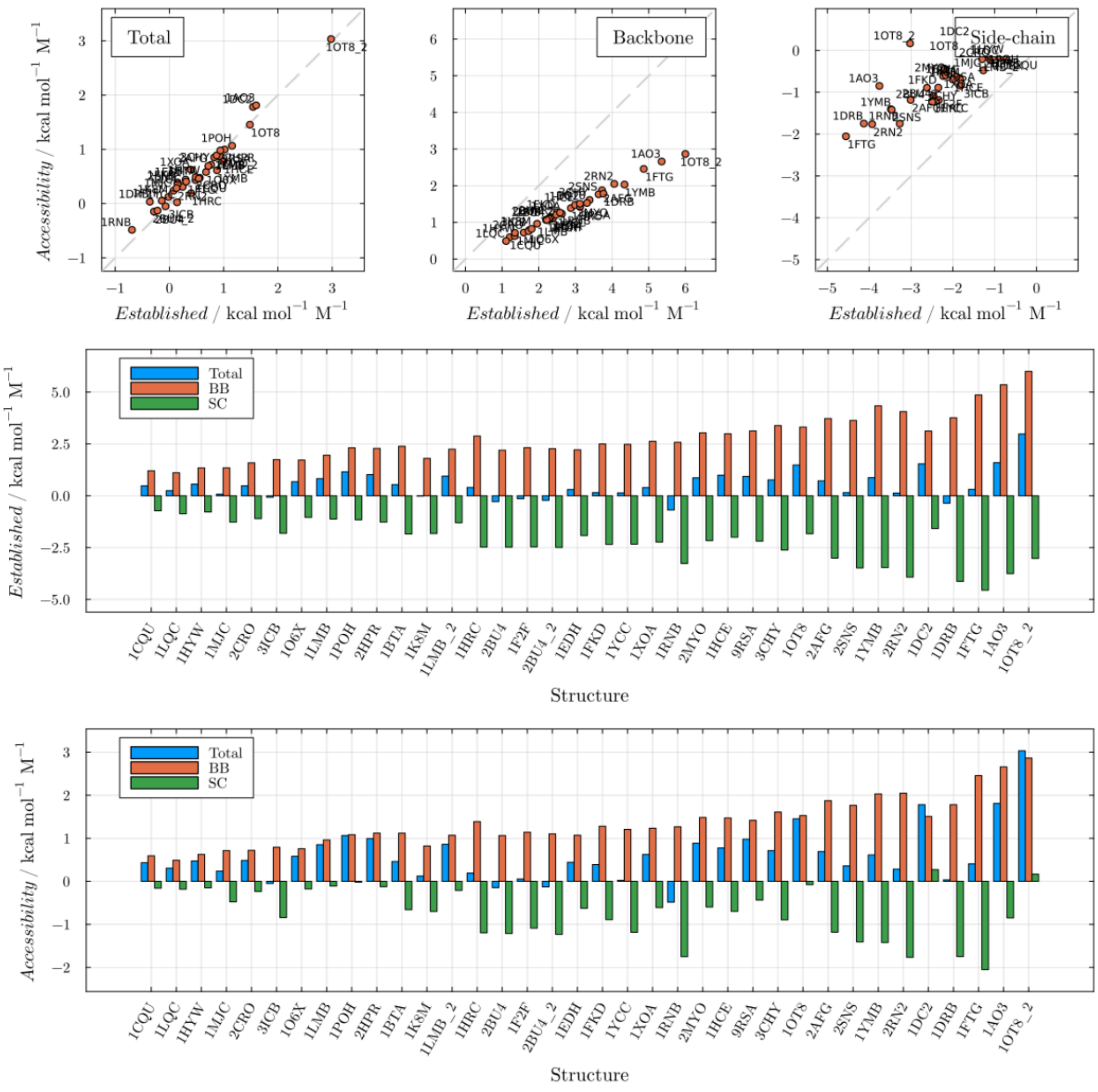


**Figure S8.** Predictions of the *Established* and *Accessibility* models for total, backbone and side-chain $m$-values in betaine.

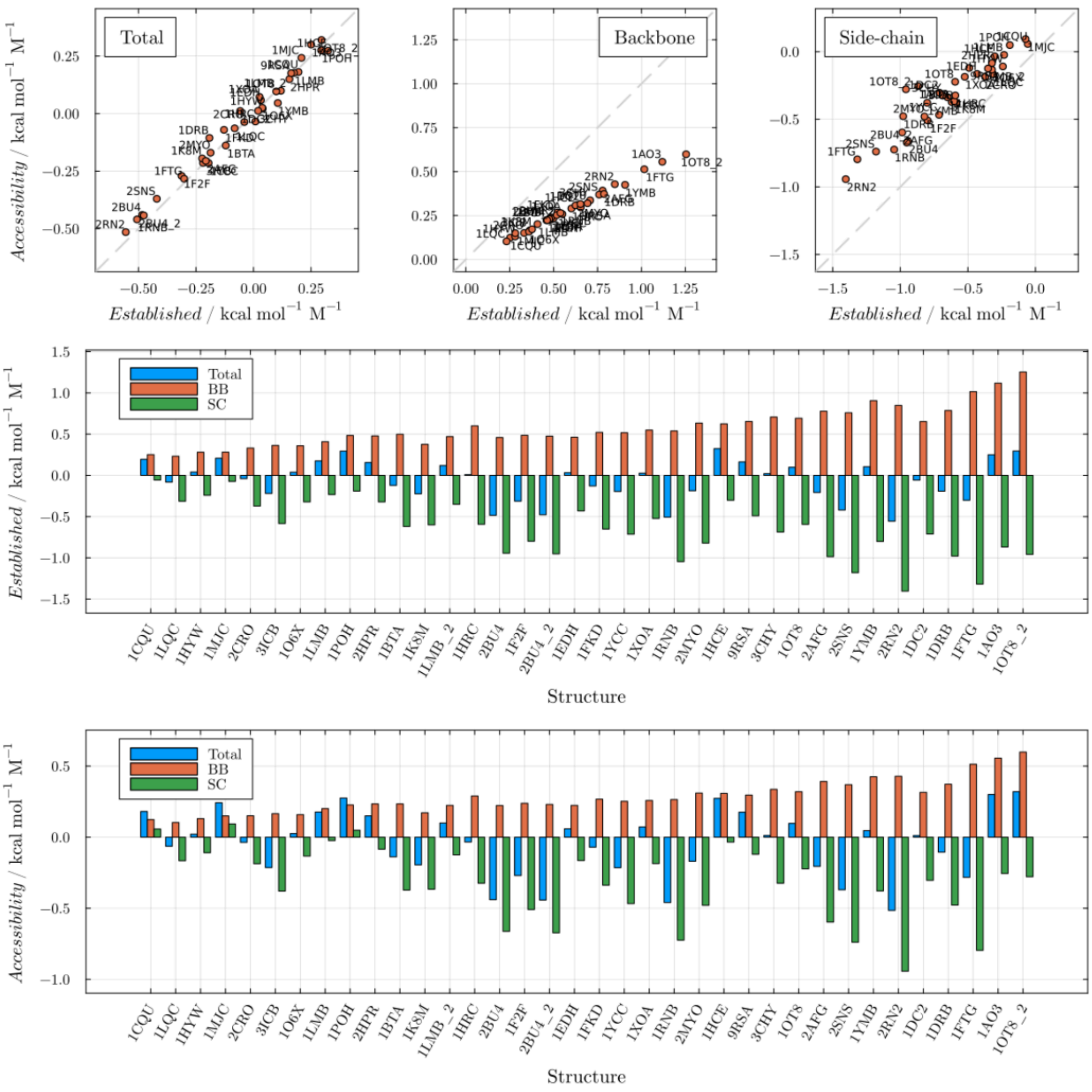


**Figure S9.** Predictions of the *Established* and *Accessibility* models for total, backbone and side-chain *m*-values in glycerol.

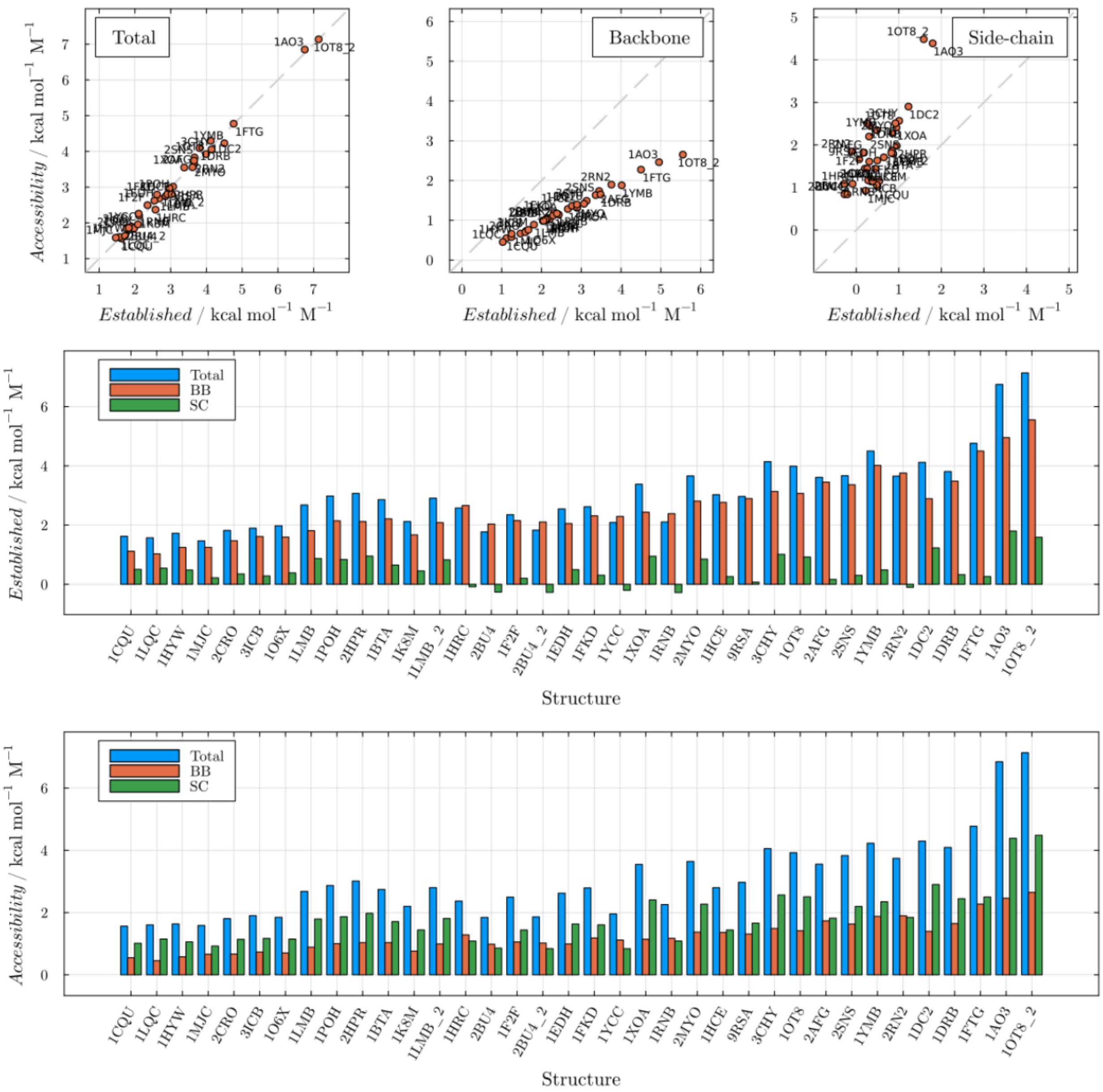


**Figure S10.** Predictions of the *Established* and *Accessibility* models for total, backbone and side-chain *m*-values in trehalose.

### S3. Predictions of *m*-values, and backbone and side-chain contributions, of the *Universal-backbone* vs *Record* models for urea.

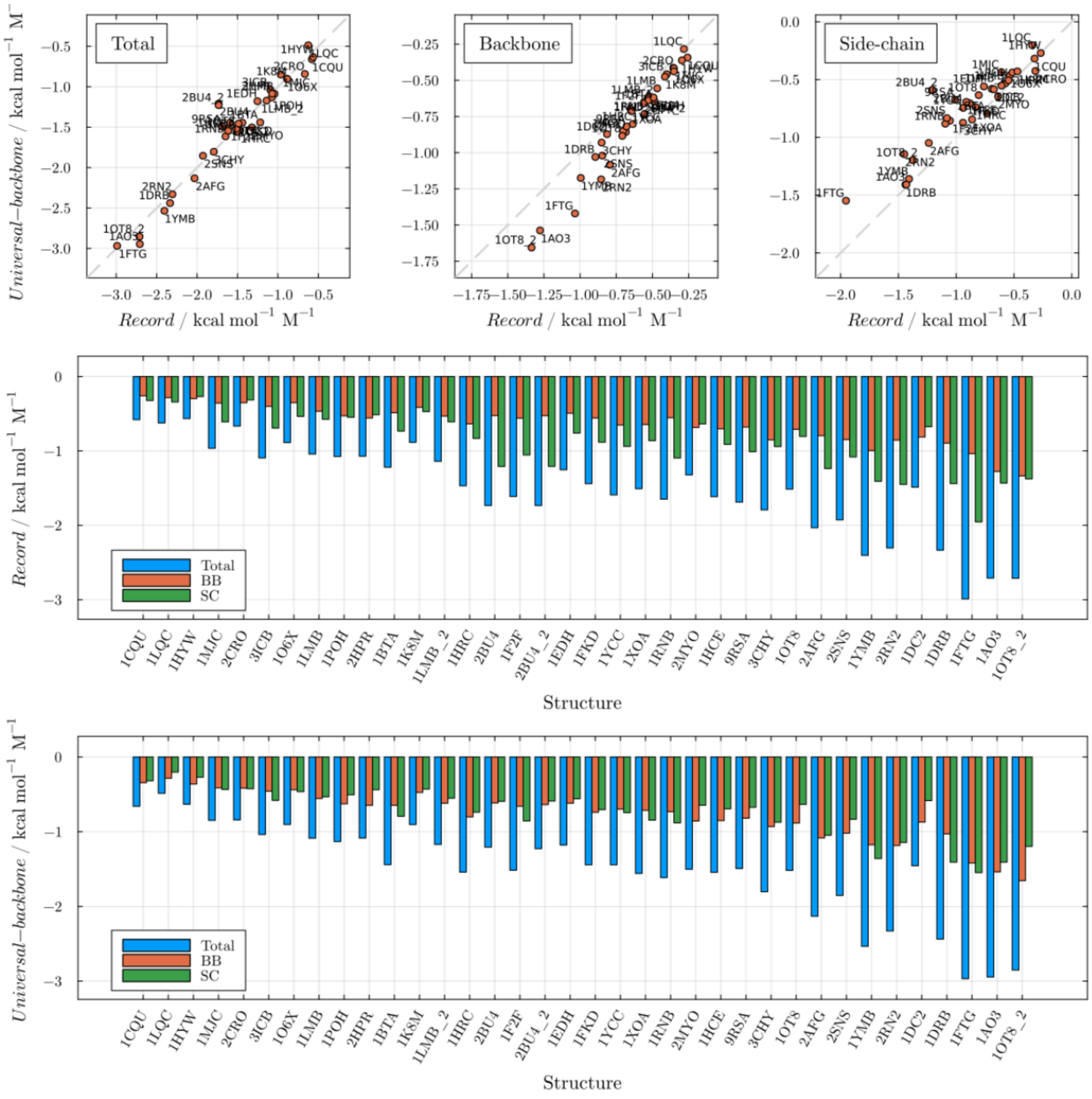


**Figure S11.** Predictions under the *Universal-backbone* and *Record* frameworks for urea support the balanced side-chain vs. backbone contributions, with the *Record* model somewhat predicting comparable but smaller backbone and greater side-chain contributions.

### S4. Predictions of *m*-values of the *Established* and *Record* models for protecting osmolytes.

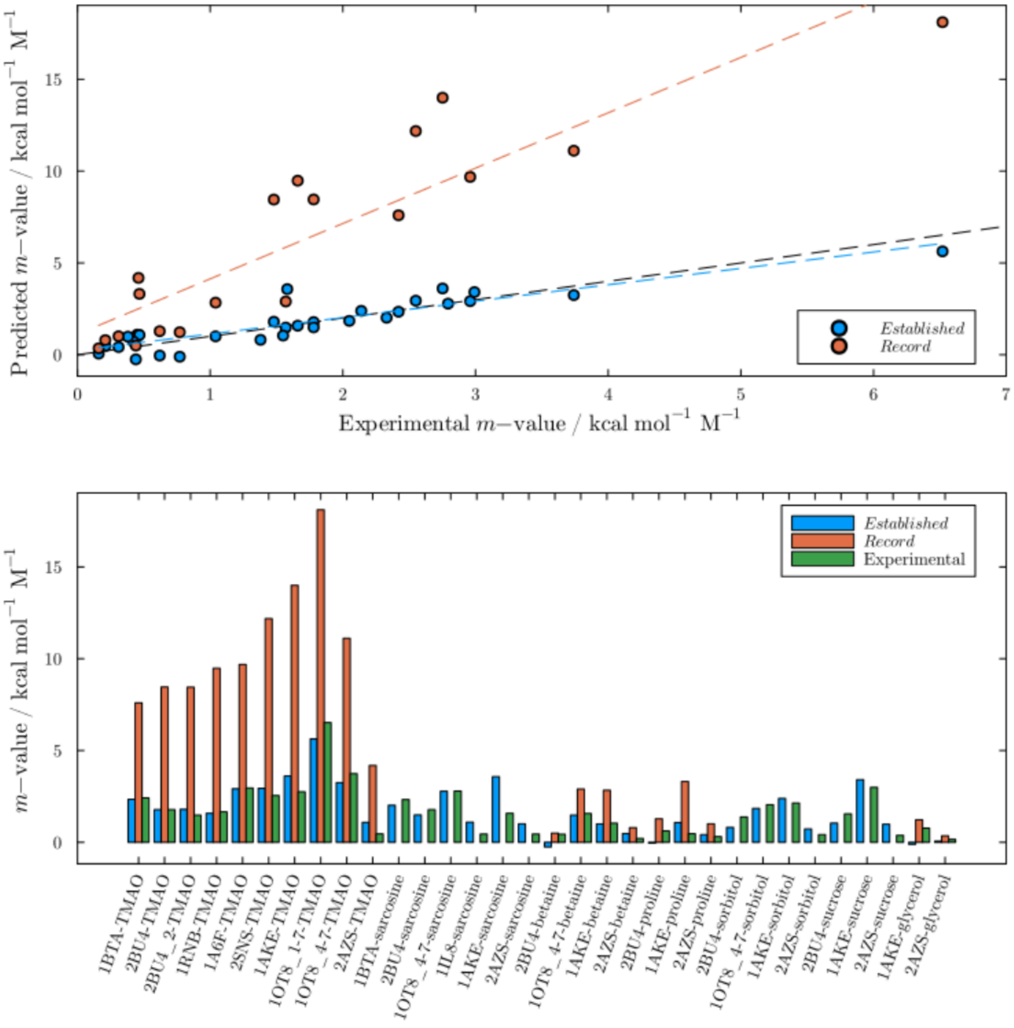


**Figure S12.** Predictions of denaturation *m*-values with the *Record* and *Established* models, for various protecting osmolytes. The denatured states considered here are an extended chain for *Record* models and the mean Creamer denatured state for the *Established* model. The *Record* model predictions for the osmolytes cannot be fitted with a single denatured state surface area model, and the overestimated values for TMAO are discussed in Section S6.

### S5. Transfer free energies of denaturation and dimer dissociation for SH3 and GB1, using the *Accessibility, Established, Universal-backbone* and *Record* models.

In this section we present the predictions of all models against the experimental data of Rydeen et al. (1), which includes denaturation free energies for the SH3 protein and dimerization free energies of the GB1 dimer, in various osmolytes.

All models are implemented in PDBTools.jl version 3.39.0 (2). The *Record* model uses parameterizations that are obtained from (3) for urea and betaine, from (4) for TMAO, from (5) for proline, from (6) for trehalose, and from (7) for glycerol. For denaturation predictions, Figure S12 considers a fully denatured state (extended chain) for *Record* models, and the mean denatured Creamer model (8) for the other models. The use of extended chain model is arguably correct for the *Record* model only for denaturation in urea, and for other osmolytes empirical fitting of the predictions is performed independently for each cosolvent. The choice of the denatured state surface area model is a fundamental limitation in the practical validation of the models, as it remains an adjustable parameter.

The purpose of Figure S13 is then to show that urea predictions are all satisfactory, but that obtaining a reassuring model for other osmolytes remains a challenge. Using the standard choice of denatured model (the mean Creamer model) for the *Accessibility*, *Established*, and *Universal-backbone* models, good predictions are obtained for the *Accessibility* and *Established* models for TMAO, trehalose, sarcosine, betaine, and sucrose, and greater deviations are observed for proline, sorbitol, and glycerol, but the directionality of the effects is always correct. The predictions of the *Universal-backbone* model are always underestimated, for the reasons discussed in the main article. The *Record* model is much more sensitive to the nature of the denatured state, and the predictions are only indicative that the model captures the correct sign of the osmolyte effects on protein denaturation.

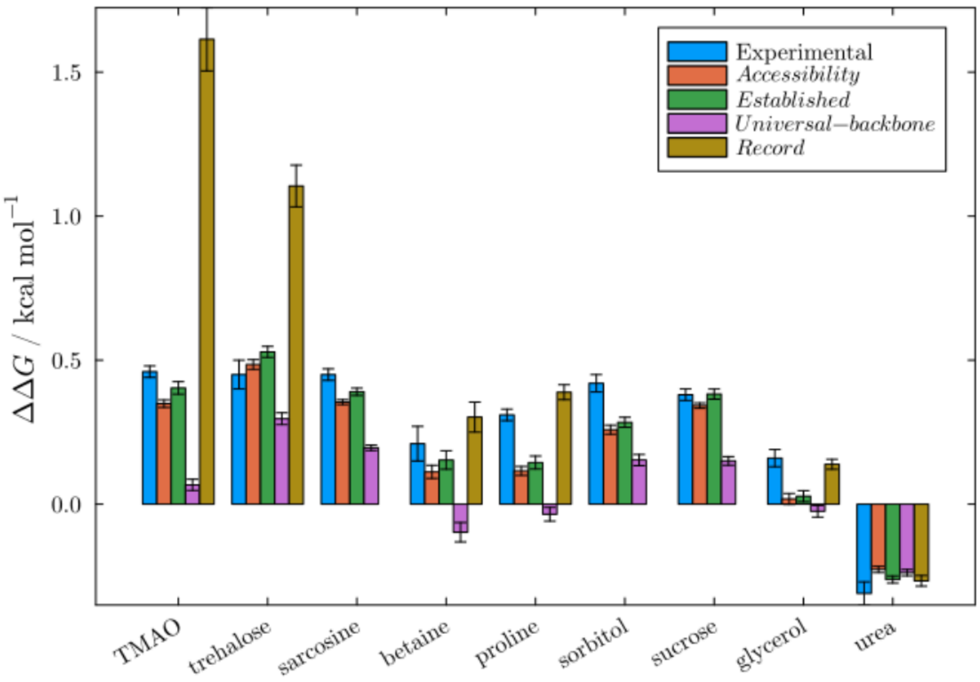


**Figure S13.** Experimental and predicted denaturation free energies for SH3 denaturation in different cosolvents. The denatured surface areas are the "average" surface area for Creamer models commonly used in combination with the *Established* model, and a fully extended chain as proposed for urea for the *Record* model. Varying the surface areas independently for each osmolyte allows fitting the data in both cases.

The predictions of the effects of osmolytes on GB1 dimer dissociation, shown in Figure S14, are more interesting to compare because the bound and dissociated structures are well defined if one assumes that the dissociated form consists of two monomers with roughly conserved monomeric folds.

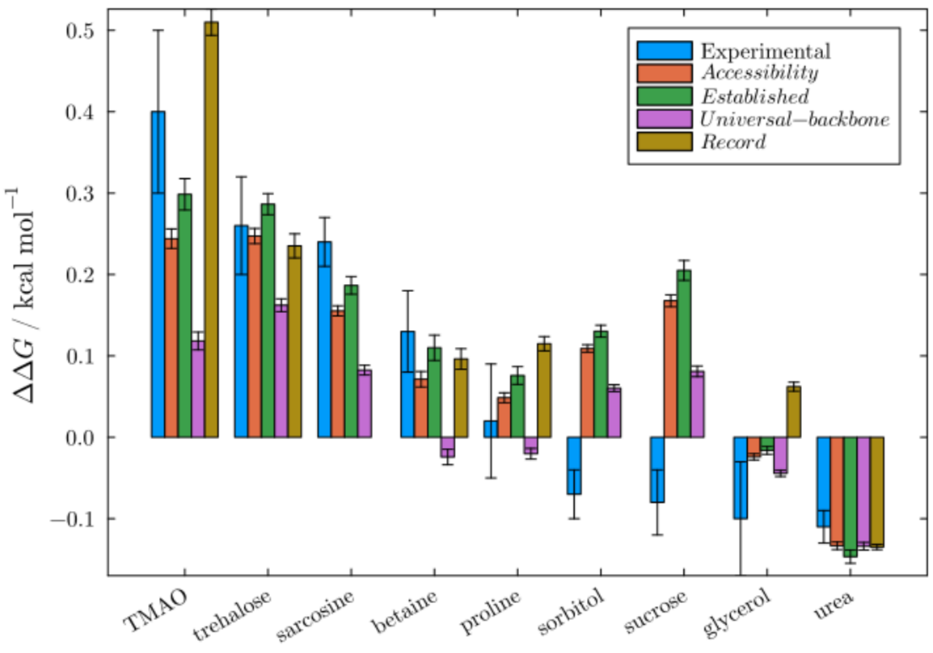


**Figure S14.** Experimental and model predictions for GB1 dimer dissociation, where the experimental data is reported by (1). Standard deviation of the experimental data and of predictions made for the 20 NMR models of the 2RMM are shown.

The predictions of each model for each osmolyte effect vary in quality but, except for the predictions in sorbitol and sucrose the *Accessibility*, *Established*, and *Record* models provide adequate qualitative predictions. In TMAO, the *Record* model overestimates the free energy, while the other models somewhat underestimate it. In urea all predictions are equally satisfactory. Notably for the purposes of the present work, the *Universal-backbone* model provides systematic deviations in all protecting osmolytes, thus indicating that a revision of the distribution of effects was effectively necessary.

### S6. *Record* model predictions for TMAO in comparison with molecular dynamics simulations

Here we discuss the predictions of the *Record* model for TMAO, parameterized as described in (4). TMAO was the reference osmolyte for the validation of the models against simulations because validated Kirkwood-Buff force-fields exist (see Supplementary Section S7). Also, TMAO is one of the most widely studied protecting osmolytes, observed to stabilize the folded and oligomeric states of proteins.

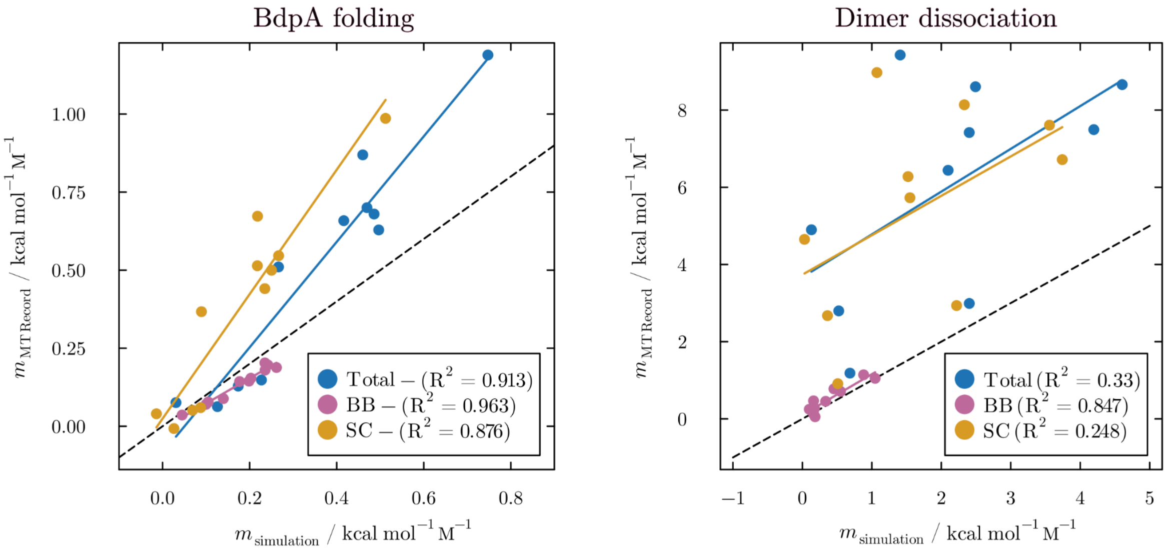


**Figure S15.** Predictions of *m*-values of unfolding and dimer dissociation using the *Record* model in comparison with simulated predictions, for TMAO. With the current available parameterization, the *Record* model overestimates significantly the side-chain contributions for the transfer free-energies, leading to overestimated total predictions. The estimates for the backbones agree nicely with the predictions from the simulations.

When comparing the model predictions against simulations, the structures are well defined and no ad-hoc assumptions are necessary concerning the initial and final accessible surface areas, thus being a nice way to validate the model predictions whenever the force-fields are appropriate.

Figure S15 shows that the total BdpA denaturation *m*-values predicted by the *Record* model are somewhat overestimated relative to the predictions from the simulations, and that the overestimation increases with the extent of denaturation. Interestingly, the predictions of the backbone contributions agree with the simulated results almost quantitatively, and the overestimated total *m*-values follow uniquely from the erroneous estimates of the contributions of side chains. The same pattern is observed in the predictions of dimer dissociation, with a near quantitative agreement of backbone contributions, but a systematic and highly overestimated contribution of side chains, leading to greater totals than what is computed from the simulations. These overestimated predictions are consistent with the TMAO predictions in Figures S12 and S13: The *Record* model provides greater TMAO osmolyte dimerization protection than what is observed experimentally or predicted by the other models.

These overestimated predictions of the *Record* model under the (4) parameterization limits the degree of interpretation that this model can provide to the important backbone vs. side-chain distribution of effects. At the same time, given the near quantitative agreement of the backbone contributions with the predictions of the simulations, and the fact that the simulation total free energies derive from important side-chain contributions, there is no apparent interpretation other than that the side chains should contribute more than what is predicted by the *Established* model. Thus, the *Record* models, with a different construction and using independent experimental data support, under reasonable assumptions, the increased side-chain relevance for the osmophobic effect.

Although beyond the purpose of the current work, it is worth speculating why the side-chain contributions for TMAO are overestimated in the *Record* model under the used parameterization. The reason seems to be that the experimental data used by (4) to parametrize the model does not include highly hydrophobic compounds. This probably limits the quality of the predictions of the groups of atoms that interact with TMAO, resulting in the observed skewed predictions for the side chains.

### S7. Molecular Dynamics Simulations Protocols

The protein-water-TMAO systems were modeled using the CHARMM36 (9) force field for the proteins (BdpA and Dimers) and the TIP3P model (10) for water. TMAO was described using the Netz force field (11) with the Shea group's scaling correction (12) to ensure the accurate reproduction of preferential binding coefficients in ternary protein/water/TMAO solutions (13). Systems were minimized by up to 20,000 Steepest Descent steps and equilibrated for 1 ns in a constant-volume and constant-temperature ensemble (NVT), followed by 1 ns of constant-temperature and constant-pressure (NPT) simulation. Production simulations were performed in GROMACS 2021.2 (14) in the NPT ensemble at 298.15 K and 1 atm.

Atomistic molecular dynamics simulations of the BdpA (PDB ID: 1BDD) (15) protein in a TMAO solution (0.5 mol $L^{-1}$) were based on the protocol described previously (16). Conformations from the BdpA folding ensemble - originally generated via $C\alpha$-Structure-Based Models ($C\alpha$-SBM) (17, 18) and reconstructed at the all-atom level using Pulchra (19) software - were solvated in orthorhombic boxes using Packmol (20, 21). To preserve the underlying SBM topology during the simulations, harmonic potentials with 25 kcal $mol^{-1}$ Å$^{-2}$ force constants were applied to the $C\alpha$ atoms.

To investigate the structure of dimeric proteins, a set of 10 protein dimers was selected from the Protein Data Bank (PDB ID: 1RPO, 2AG8, 2F3M, 2ILK, 2RMM, 3E2D, 3ENJ, 3GRS, 3PZA and 3REM) (22–31) based on their structural characteristics. Each dimer was placed in a cubic box and solvated under the corresponding solution conditions (water and TMAO 1.0 M solutions). After energy minimization, the systems were subjected to 5 ns of NVT equilibration. Subsequently, 10 ns of NPT equilibration was performed to constant temperature and pressure. To preserve the experimentally observed dimeric architecture throughout the simulations, positions restraints were applied to the C$\alpha$ atoms of the protein, with a force constant of 10 kcal $mol^{-1}$ $Å^{-2}$. All simulations were performed using the leap-frog integration algorithm with a 2 fs integration time step. Bonds involving hydrogen atoms were constrained using the LINCS algorithm(32). Short-range electrostatic and van der Waals interactions were calculated using a 1.0 nm cutoff. Long-range electrostatic interactions were treated using the Particle Mesh Ewald (PME)(33, 34) method. Temperature was controlled using the V-rescale, with the protein and non-protein components coupled separately using a coupling time constant of 0.1 ps and a reference temperature of 298.15 K for both groups. During the production simulations, isotropic pressure coupling was applied using the C-rescale barostat, with a reference pressure of 1 bar and a pressure coupling time constant of 2.0 ps. Molecular dynamics simulations were performed for 100 ns for each system. The same simulation protocol was consistently applied to all dimers and solution conditions.

Transfer free energies were obtained from the simulations using

$$\mathrm{TFE} = \frac{RT}{1 - c_{os}(G_{WO} - G_{OO})}(G_{PW} - G_{PO})$$

where $G_{PW}$ and $G_{PO}$ are the protein-water and protein-cosolvent Kirkwood-Buff (KB) integrals (35, 36). The first term depends on the properties of the pure solvent, where $c_{os}$ is the concentration and $G_{WO}$ and $G_{OO}$ the water-cosolvent and cosolvent-cosolvent KB integrals. The $m$-values follow from the difference of TFEs of two states (native vs. denatured or dissociated). Backbone and side-chain contributions can be estimated, for example, by decomposing the KB integrals into contributions of the protein atoms closest to each region in space when integrating density fluctuations, in a Voronoi scheme similar to that of Best and co-workers (37). We apply the same strategy, with the local solvent density associated with the minimum distance to any solvent molecule atom, following the minimum-distance distribution framework (38). Thus, hydrogen-bonding interactions essentially always determine which protein atom mandates the local accumulation of a solvent molecule, making this a better proxy for the effect of local interactions on solvent organization. All calculations were performed with the ComplexMixtures.jl package (39), version 2.18.0, for which the spatial decomposition of KB integrals was implemented under the :kbi option of the *contributions* function.

### S8. Model predictions in selected dimers

The protein dimer structures proposed for the $m$-value comparison are selected based on available biophysical protein experiments described in the literature. These are the Vibrio alkaline phosphatase (PDB ID: 3E2D) (40, 41), human glutathione reductase (PDB ID: 3GRS) (42, 43), pig heart citrate synthase (PDB ID: 3ENJ) (44, 45), human glutathione S-transferase M1a-1a (PDB ID: 2F3M) (24, 46, 47), human interleukin-10 (PDB ID: 2ILK) (48), the Escherichia coli Rop protein (PDB ID: 1RPO) (49, 50), Neisseria meningitidis pyrroline-5-carboxylate reductase (PDB ID: 2AG8) (23), Pyrococcus furiosus rubrerythrin (PDB ID: 3PZA) (51), Pseudomonas aeruginosa isochorismate-pyruvate lyase (PDB ID: 3REM) (52), and the engineered GB1-A34F dimer (PDB ID: 2RMM) (1, 26).

The selected structures were obtained from the Protein Data Bank as biological assemblies assigned by the authors as dimers. These assemblies were used as the associated states, whereas the two constituent monomers separated from each assembly were treated as the dissociated states. At atom sites with multiple conformations, the conformation with the highest occupancy was retained. Modified residues were excluded from the *m*-value calculations. Dissociation *m*-values were computed using the transfer models implemented in the PDBTools.jl package (https://m3g.github.io/PDBTools.jl - version 3.39.0) (2), allowing direct comparison between the *Established, Universal, Record* and *Accessibility* model predictions for cosolvents urea, betaine, proline, glycerol, sorbitol, sucrose, trehalose, sarcosine, and TMAO.

### S9. Shielding of urea-backbone hydrogen bonds by side chains

Tripeptides with the sequence Gly-X-Gly (GXG), where X represents the target amino acid, were constructed using AmberTools (53) with either neutral ($NH_2$/COOH) or charged ($NH_3^+$/$COO^-$), or capped (Ace/$NH_2$) terminal groups. Each peptide was placed in a cubic box with a side length of 50 Å and solvated in a 1.0 M aqueous urea solution using Packmol (20, 21). Interactions were modeled using the KBFF force field for urea (54) and peptides (55), combined with the SPC/E water model (56). Following the equilibration protocol described previously for BdpA, production simulations were carried out in the NPT ensemble at 298.15 K and 1 atm for 200 ns without positional restraints on the peptides.

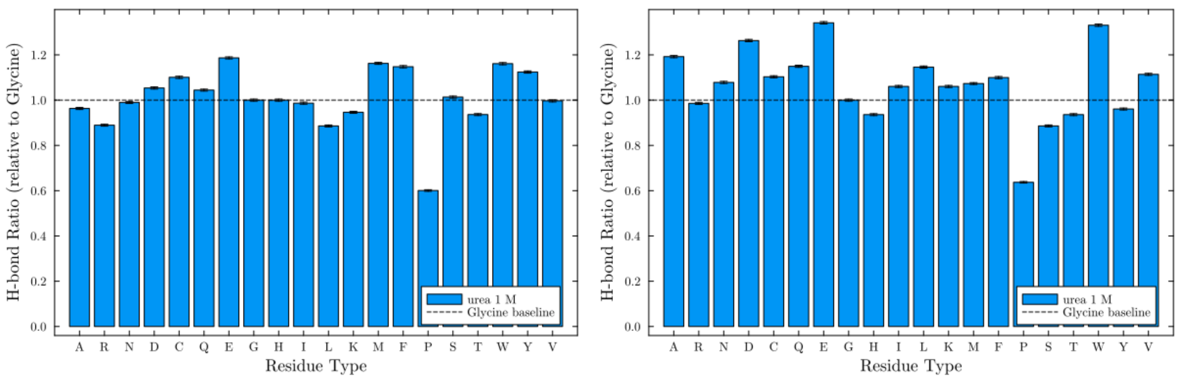


**Figure S16.** Number of hydrogen bonds of the backbone of amino acid residues with urea, in tri-peptides GXG, where X is a different amino acid type, relative to Glycine. The side chains generally do not promote a significant shielding of hydrogen bond attack by urea. The images refer to tri-peptides with (left) charged terminal groups and (right) neutral terminal groups.

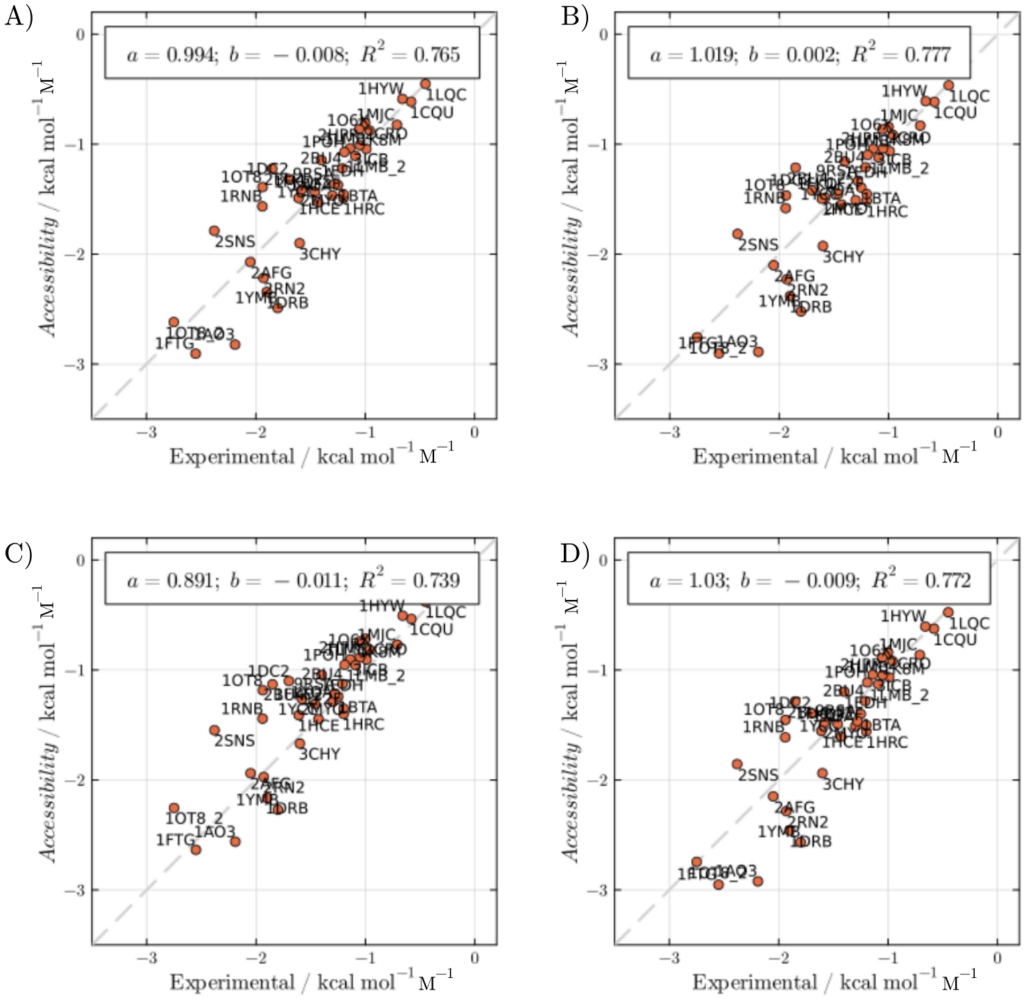


**Figure S17.** Correlations of urea *m*-values computed with the *Accessibility* model with experimental data, using different backbone accessibilities based on hydrogen-bond ratios relative to Glycine. A) fully accessible (all 1.0), and from simulations of tri-peptides with B) charged terminals, C) neutral terminals, and D) capped terminals.
